# A novel pathosystem between *Aeschynomene evenia* and *Aphanomyces euteiches* reveals new immune components in quantitative legume root-rot resistance

**DOI:** 10.64898/2026.01.11.698850

**Authors:** Madeleine Baker, Yves Martinez, Jean Keller, Baptiste Sarrette, Marjorie Pervent, Cyril Libourel, Aurélie Le Ru, Maxime Bonhomme, Clare Gough, Baptiste Castel, Jean-François Arrighi, Christophe Jacquet

## Abstract

Legumes are pivotal for sustainable agriculture, yet their productivity is hindered by soilborne pathogens such as *Aphanomyces euteiches*. This study introduces *Aeschynomene evenia* as a novel model to investigate legume immunity and its interplay with Nod factor-independent symbiosis (NFIS).

Inoculation of *A. evenia* accessions with *A. euteiches* strains revealed a range of quantitative resistance levels. Phenotypic and cytological analyses showed a compatible interaction, including root browning, intracellular colonisation, and oospore production; along with partial resistance traits such as phenolic accumulation at the endodermal barrier.

Transcriptomic analyses at 1 and 3 dpi identified 3,403 differentially expressed genes (DEGs), with upregulated genes associated with pathogen perception, immune signalling and specialised metabolite biosynthesis. Kinase-mediated signalling, ROS homeostasis, and WRKY transcription factors were among the most enriched functional categories. Comparative transcriptomics with *Medicago truncatula* confirmed conserved immune responses, while *A. evenia* displayed broader transcriptional repression.

Integration with symbiotic transcriptomic data revealed overlapping gene signatures, including *AeRLCK2* and *AeCRK*, previously found to be NFIS-specific, as putative dual-function kinases. *Aerlck2* and *Aecrk* mutants exhibited enhanced resistance to *A. euteiches*, supporting their roles in immune regulation.

This work positions *A. evenia–A. euteiches* as a valuable system to dissect legume quantitative resistance and its intersection with symbiosis.

## 1 Introduction

Legume crops play a pivotal role in global agriculture due to their ability to fix atmospheric nitrogen through symbiotic interactions with rhizobia, significantly reducing dependency on synthetic fertilizers and enhancing sustainability in diverse farming systems (Herridge *et al*., 2008; Kebede, 2021).

In a comprehensive assessment of crop losses due to pathogens, Savary *et al*. (2019) highlighted that legumes, including soybean, suffer significant yield reductions due to fungal, bacterial, and viral diseases. These losses not only threaten food security but also undermine the environmental benefits provided by legumes. Among important crop and forage legumes, such as pea (*Pisum sativum*) and alfalfa (*Medicago sativa*), several soil-borne pathogens like *Aphanomyces euteiches* (Kumar *et al*., 2021), *Fusarium* spp. (Achari *et al*., 2021), *Phytophthora medicaginis*, and *Pythium* spp. (Larsen *et al*., 2023) are of particular concern. These pathogens cause root rot diseases and seedling damping-off, leading to severe yield and quality losses under conducive conditions.

Among them, *Aphanomyces euteiches* stands out as a major soil-borne threat to legumes due to its broad host range, including pea, alfalfa, clover, lentil, bean, vetch, and chickpea (Gaulin *et al*., 2007; Moussart *et al*., 2008). Its long-lived oospores enable persistence in agricultural soils, making it one of the most difficult pathogens to manage (Gaulin *et al*., 2007). Since no effective treatments are available to control the disease, identifying new sources of resistance and uncovering key genes involved in plant immunity against *A. euteiches* remain crucial challenges.

Over the years, genetic studies have been instrumental in identifying loci and genes involved in resistance to *Aphanomyces euteiches*. In pea (*Pisum sativum*), early studies led to the discovery of quantitative trait loci (QTL) associated with partial resistance to the pathogen (Pilet-Nayel *et al*., 2002; Lavaud *et al*., 2015) and more recent advances have further expanded this knowledge (Lavaud *et al*., 2024). In parallel, in the model legume *Medicago truncatula*, QTL mapping and genome-wide association studies (GWAS) have identified several loci linked to resistance to *A. euteiches* (Bonhomme *et al*., 2014 and 2019). These approaches have provided a solid foundation for understanding the genetic architecture of resistance to this oomycete in legumes.

A complementary strategy to identify genes involved in plant immune responses to *A. euteiches* has relied on functional analyses using RNA interference (RNAi) or mutant approaches. Candidate genes were either identified through omics-based analyses (Kiirika *et al*., 2012; Yadav *et al*., 2019) or derived from mutants previously characterized for roles in symbiotic processes, suggesting crosstalk between symbiosis and immunity, particularly against *A. euteiches* (Rey and Jacquet, 2018). This latter approach in *M. truncatula* has been particularly successful in identifying membrane receptors that play a dual role in symbiosis and immunity, including MtNFP (Rey *et al*., 2013), MtLYK9 (Gibelin-Viala *et al*., 2019), and more recently MtSOBIR1 (Sarrette *et al*., 2025).

While these findings in *P. sativum* and *M. truncatula* provide valuable insights, they primarily focus on legumes within the Inverted Repeat-Lacking Clade (IRLC) (Choi *et al*., 2019). The extent and nature of this symbiosis-immunity interaction in other legume species, especially outside of this clade, remain largely unexplored.

To address this gap, the Dalbergioid legume *Aeschynomene evenia* has emerged as a promising model. Unlike *M. truncatula*, this legume establishes symbiosis with photosynthetic *Bradyrhizobium* spp. via a Nod-Factor (NF)-independent mechanism, providing a distinct framework for studying alternative symbiotic pathways (Arrighi *et al*., 2012; Quilbé *et al*., 2021). Recent studies using EMS mutagenesis in *A. evenia* have identified key symbiotic genes, previously described in NF-dependent legumes, such as *AeNSP2* and *AeNIN*, which are also required for NF-independent symbiosis (Quilbé *et al*., 2021). In addition, novel genes potentially specific to NF-independent symbiosis, notably those encoding for the two kinases AeCRK and AeRLCK2, have been identified (Quilbé *et al*., 2022; Horta Araújo *et al*., 2025). These findings highlight the unique potential of *A. evenia* to uncover new molecular players involved in plant-microbe interactions. However, there is currently no established pathosystem to fully explore both the immune mechanisms of *A. evenia* and their potential crosstalk with NF-independent symbiosis (NFIS).

In this study, we leverage the compatibility of the interaction between *A. evenia* and *A. euteiches* to establish a novel pathosystem. Through transcriptomic and cytological analyses, together with the examination of natural variation across multiple *A. evenia* accessions, we provide the first evidence for partial and quantitative resistance in *A. evenia* against this oomycete. Comparative transcriptomics across species revealed a common response to *A. euteiches* between *A. evenia* and *M. truncatula*, and within *A. evenia* we identified a core set of genes responding to both pathogenic (*A. euteiches*) and symbiotic (*Bradyrhizobium* spp.) interactions.

Finally, analyses of symbiotic mutants uncovered AeRLCK2 and AeCRK as negative regulators of resistance to *A. euteiches*, underscoring the potential of this system to yield new insights into legume immunity and its intersections with symbiotic signalling.

## 2 Methods

### 2.1 Plant Material and Growth Conditions

The *Aeschynomene evenia* natural accessions (Brottier *et al*., 2018; Chaintreuil *et al*., 2018) used in this study included the sequenced reference accession CIAT 22838 (n°76, Malawi, East Africa), CIAT 08251 (n°79, Brazil, South America), CIAT 08254 (n°77, Brazil, South America), CIAT 08232 (n°80, Brazil, South America), and PI 225551 (n°21, Zambia, Southeast Africa).

Mutants were previously isolated from a large-scale EMS-mutagenesis screen on the CIAT 22838 (n°76) line, as described by Quilbé *et al*. (2021). Each mutant used in this study was backcrossed once. For *AeRLCK2* (<u>Ae01g26600</u>) and *AeCRK* (*<u>Ae05g12380</u>*) two independent alleles were analysed (*rlck2-1* and *rlck2-2*; Horta Araújo *et al*., 2025, *crk-1 and crk-2;* Quilbé *et al.,* 2021).

Seeds were scarified with 98% sulfuric acid for 40 min, rinsed five times with sterile distilled water, and left overnight in the dark at 22°C. Following rehydration, seeds were germinated on 1% agar plates at 25°C in the dark. Germinated seeds were transferred to 12 × 12 cm Petri dishes containing M medium, pH 5.5 (Bécard and Fortin, 1988), which is classically used to inoculate *M. truncatula* with *A. euteiches,* and placed vertically in growth chambers at 25/22°C (16 h light / 8 h dark cycle).

### 2.2 Microscopy and Image Analysis

Roots and stems were prepared for microscopy by embedding in 7% low-melting-point agarose, and 100 µm transverse sections were generated using a Leica VT1000S vibratome. Sections were stained with Wheat Germ Agglutinin-Alexa Fluor 488 conjugate (WGA-AF488) diluted in 1× PBS to visualize *Aphanomyces euteiches* mycelium. Samples were mounted in distilled water and observed using a Nikon Eclipse Ti inverted microscope equipped with a 10×/0.3 objective. A GFP filter set was used for WGA-stained mycelium (ex: 472/30 nm, em: 520/535 nm) and a DAPI long-pass filter set for plant autofluorescence (ex:390/40, em: 409 LP).

Hyphal density was quantified using ImageJ.js v0.5.7 (Schneider *et al*., 2012). The cortex surface area and the area of WGA fluorescence were measured using hue and saturation thresholds, and the percentage of colonised cortex was calculated for each plant (https://github.com/Madeleine-Baker/Aeschynomene_evenia_Baker_et_al_2025.git : Macro_root_colonisation.txt, **Supp. data: Table S1, Methods S1**). For each line, 10 to 15 plants were sectioned multiple times and analysed randomly. Statistical analyses were performed using RStudio v1.3.1093 employing Wilcoxon tests with Bonferroni correction.

For scanning electron microscopy, root samples were fixed in a 0.05M solution of sodium cacodylate buffer (pH 7.2) containing 2.5% glutaraldehyde, dehydrated in an ethanol series, and critical-point dried with liquid CO_2_. Samples were mounted with conductive silver paint on stubs and sputter-coated with platinum. Images were acquired with a Quanta 250 FEG (FEI) scanning electron microscope at 5 kV, spot size 3 and a working distance of 10 mm.

### 2.3 *Aphanomyces euteiches* culture, inoculation and symptom phenotyping

*Aphanomyces euteiches* strains ATCC201684 and RB84 (Bonhomme *et al*., 2019) were cultured on corn meal agar (17 g/L, Sigma). Zoospore inoculum was prepared by transferring agar plugs colonised by mycelium into peptone-glucose liquid medium, followed by incubation for 3 days at 20°C. After incubation, the plugs were washed six times with 100 mL of Volvic water and incubated overnight in 30 mL Volvic water to induce starvation-based sporulation. Zoospores were then harvested and their concentration was adjusted to 100,000 spores/mL using a Fuchs-Rosenthal haemocytometer. Inoculation was performed by applying 5 µL droplets of this suspension onto the root hair zone of 1-day-old seedlings.

To monitor root browning symptoms, quantitative image analysis of root phenotypes was performed at 3, 5, 7, 10, and 12 dpi using ImageJ.js v0.5.7 (Schneider *et al*., 2012). Total root surface and browning areas were measured based on hue and saturation thresholds and the resulting values were used to calculate the percentage of root browning for each plant (https://github.com/Madeleine-Baker/Aeschynomene_evenia_Baker_et_al_2025.git, and **(Supp. data; Tables S2 and S3**). For each line, 10–15 plants were analysed in three independent repeats. Statistical analyses and plots were performed using RStudio v1.3.1093 along with the packages data.table v1.15.0 (Barett *et al*., 2024), ggplot2 v3.5.0 (Wickham, 2016) and dplyr v1.1.4 (Wickham *et al.,* 2014). Pairwise Wilcoxon tests with Bonferroni correction were used for fresh weight and area under the curve of percentage of root browning across natural accessions. For mutant lines, the area under the curve of percentage of root browning was compared to the WT (n°76) using a general linear model considering the replicate effect as no interaction between replicate and genotype was observed (**Supp. data; Methods S2**).

### 2.4 RNA-seq sampling and analyses

Plants were grown on M medium for 6 days prior to inoculation. Four biological replicates were collected for each time point and condition. Inoculation was performed by applying 5 µL droplets of sterile Volvic water, either alone or containing *A. euteiches* RB84 zoospores (100,000 spores/mL), at 0.5 cm intervals along the roots. Roots from five to ten plants, either inoculated or mock-treated, were collected for each replicate at one- and three-days post-inoculation (dpi) and immediately flash-frozen in liquid nitrogen. Total RNA was extracted using the SV Total RNA Isolation System (Promega), and quantified with a NanoDrop ND-1000 spectrophotometer (Gully *et al*., 2018). mRNA libraries were prepared from non-inoculated roots and inoculated roots, collected at 1 and 3 dpi, and sequenced on an Illumina NovaSeq 6000 platform, generating 150pb oriented paired-end reads (GeT-PlaGe platform, Toulouse, France; https://get.genotoul.fr/la-plateforme/get-plage/). (SRA: SRP446020).

### 2.5 Differential gene expression analysis

Differentially expressed genes (DEGs) were identified from *A. evenia* roots under control and inoculated conditions at 1 and 3 dpi (**Supp. data; Table S4**), as well as in *M. truncatula* (genotype A17) challenged with *A. euteiches* (strain RB84) at 3 dpi (**Supp. data; Table S5**). To investigate the interplay between immunity and root nodule symbiosis in *A. evenia*, we also re-analysed RNA-seq reads from a published dataset (PRJNA399681; 10.1038/s41598-018-29301-0) using the same analysis pipeline as for the immune response (**Supp. data; Table S6**).

RNA-seq reads were processed using the Nextflow v23.10.0 (Tommaso *et al*., 2017) pipeline NF-CORE/RNA-seq v3.14 (Ewels *et al*., 2020), with the star_salmon option to align and quantify reads as well as ‘-nextseq 30 --length 50’ as extra parameters of TrimGalore v0.6.7 (Krueger *et al*., 2021) to remove reads with quality lower than 30 or a length shorter than 50bp. Ribosomal RNA was also removed through the “-remove_ribo” option of SortMeRNA v4.3.4 (Kopylova *et al*., 2012). The pipeline was run with the GenoToul configuration (https://github.com/nfcore/configs/blob/master/docs/genotoul.md) and used the following softwares and languages: bedtools v2.30.0 (Quinlan and Hall, 2010), R v4.0.3, v4.1.1 and v4.2.1 (R Core Team, 2018), DESeq2 v1.28.0 (Love *et al*., 2014); dupradar v1.28.0 (Sayols *et al*., 2016), fastQC v 0.12.1 (Andrews, 2010), fq v0.9.1 (https://github.com/stjude-rust-labs/fq), gffread v0.12.1 (Pertea, 2020), Perl v5.26.2 (Wall *et al*., 2000), Python v3.9.5 (Van Rossum, 2009), RSEM v1.3.1 (Li and Dewey, 2011), STAR v2.7.10a (Dobin *et al*., 2013), Picard v3.0.0 (Picard toolkit, 2019), Qualimap v2.3 (Okonechnikov *et al*., 2016), RseQC v5.02 (Wang *et al*., 2012), Salmon v1.10.1 (Patro *et al*., 2017), Summarized Experiment v1.24.0, (https://bioconductor.org/packages/release/bioc/html/SummarizedExperiment.html), Samtools v1.16.1 (Danecek *et al*., 2021), StringTie v2.2.1 (Kovaka *et al*., 2019), tximeta v1.12.0 (Love *et al*., 2020), UCSC v377 and v445 (https://github.com/ucscGenomeBrowser/kent).

Finally, DEGs were independently estimated for each time point using edgeR v4.4.1 (Chen *et al*., 2024). Briefly, genes with low variance across conditions were removed using the default edgeR ‘filterByExpr’ function followed by normalisation and DEG identification using the quasi-likelihood method with a false discovery rate (FDR) threshold of 0.05 (**Supp. data; Method S3; Table S4 and S5**).

### 2.6 Functional annotation and enrichment

To perform functional enrichment of DEGs, predicted proteins of *A. evenia* and *M. truncatula* were first subjected to functional annotation using the InterProScan suite v5.64-96.0 (Jones *et al.,* 2014) with default parameters and the ‘-goterms’ and ‘-iprlookup’ options enabled. Functional enrichment of GO and IPR terms was carried out using clusterProfiler v4.12.6 (Wu *et al*., 2021) with default parameters (**Supp. data; Method S3; Tables S7 and S8**), considering DEGs with |logFC| > 1 as the foreground set. The same approach was applied on commonly differentially expressed genes in response to *Bradyrhizobium spp. A. euteiches* (**Supp. data; Tables S9 and S10**).

### 2.7 GO and InterPro domain enrichment figures

From the GO term enrichment analysis, significantly enriched terms were classified by the highest gene ratio. Bubble plots were generated using the SRplot enrichment bubble plot tool (Tang *et al*., 2023).

### 2.8 Comparative transcriptomics immunity versus symbiosis

#### 2.8.1 Heatmap

All significant DEGs in each condition were combined, producing a list of genes that are significantly deregulated at any time point in response to *A. euteiches* and/or *Bradyrhizobium spp.* and plotted using the Heatmap function from the ComplexHeatmap package (Gu *et al*., 2016). Differentially expressed genes were categorised by profile, identifying 63 genes among significant DEGs in response to *Bradyrhizobium spp.* inoculation that presented mixed expression profile across the different time points and were classified as “other”. Genes that were upregulated at least once in each response were classified as “common up” and similarly for “common down” for downregulated genes. Opposite expression profile genes were identified as those upregulated once in response to *Bradyrhizobium spp.* and downregulated at least once in response to *A. euteiches* and vice versa (**Supp data; Method S4 and Table S6**).

#### 2.8.2 GO enrichment analysis

Enrichment analysis was then performed on each category of commonly and opposite deregulated genes, using the method previously described in section 2.6. The genes analysed excluded genes with mixed profiles, representing 8/215 genes among common up, 14/517 among common down and none among opposite-profile genes.

### 2.9 Orthogroups reconstruction

To identify the shared and specific transcriptomic responses of *A. evenia* and *M. truncatula* to *A. euteiches,* orthogroups were reconstructed using OrthoFinder v2.5.5 (Emms and Kelly, 2019) with the ‘diamond_ultra_sens’ and ‘-M msa’ options enabled. For robust orthogroup reconstruction, 15 Fabaceae species covering the main taxonomic orders were included (**Supp data; Table S11**).

### 2.10 Orthogroup comparison

In order to compare transcriptomic responses of *A. evenia* and *M. truncatula*, orthogroups were examined to determine the percentage of up- or downregulated genes for each species belonging to each orthogroup (**Supp data; Table S12**). All orthogroups containing upregulated genes for each species were extracted, the same approach was applied to orthogroups containing downregulated genes (it is possible for orthogroups to be in both up and down regulated categories due to containing up- and down-regulated genes) (**Supp data; Tables S13 and S14**). Venn diagrams were generated to represent species-specific and shared differentially regulated orthogroups in *A. evenia* and *M. truncatula* using Jvenn (Bardou *et al*., 2014).

A GO-term enrichment analysis was then performed on orthogroups containing significantly deregulated genes in both species. To do so, commonly up- or downregulated orthogroups were filtered, and the GO terms associated with each of these genes extracted (**Supp data; Table S15**). In parallel, all orthogroups containing expressed genes in both species were filtered separately, and GO terms associated to each genes merged to construct the reference universe. Enrichment of GO terms associated to up- and downregulated genes was then performed as described above using clusterProfiler, with the GO terms of expressed genes as background (**Supp data; Table S16**).

## 3 Results

### 3.1. Aeschynomene evenia is a host to Aphanomyces euteiches

After inoculation with *A. euteiches* zoospores of strains ATCC201684 and RB84, plants of the *Aeschynomene evenia* reference n°76 genotype exhibited clear and reproducible symptoms, including root browning, altered root architecture and overall stunted growth. The severity of these symptoms varied depending on the strain of *A. euteiches* used for inoculation (**Fig. 1**).

**Figure 1:**
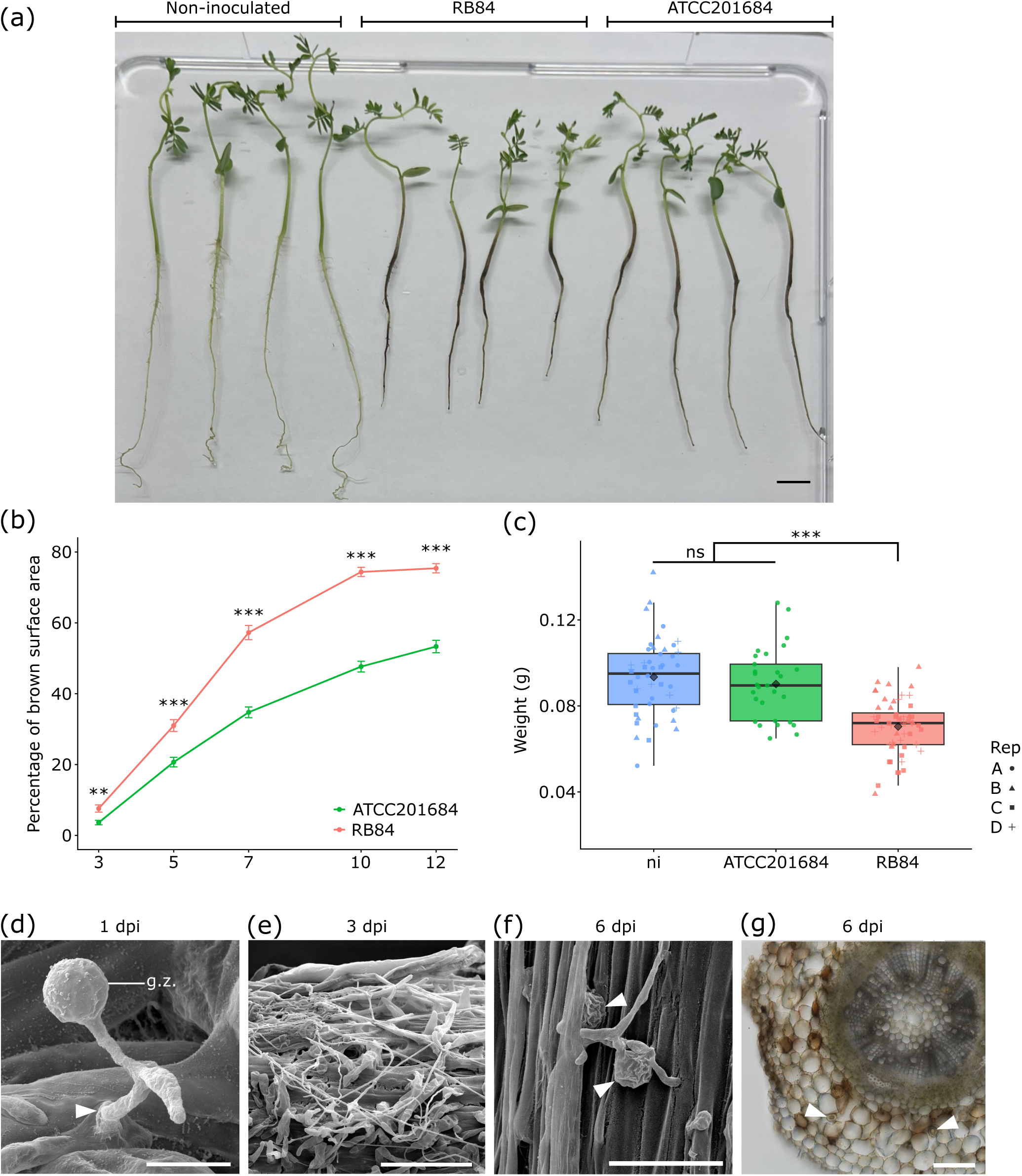
The reference n°76 line of *Aeschynomene evenia* is a host to *Aphanomyces euteiches*. **(a)** Phenotypes of non-inoculated and inoculated n°76 plants at 21 days post inoculation (dpi) with two *A. euteiches* strains (ATCC201684 and RB84). Root browning and developmental impact are associated with pathogen colonisation. Scale bar 1 cm. **(b)** Average percentage of root surface browning at 3, 5, 7, 10 and 12 dpi (N = 40-45 plants). Comparisons of browning induced by each of the two strains, at each time-point. ** p-value < 0.01; *** p-value < 0.001 (Wilcoxon test with Bonferroni correction), error bars represent standard error. **(c)** Fresh weight of non-inoculated (ni) and inoculated (ATCC201684 and RB84) n°76 plants at 21 dpi. *** p-value < 0.001 (pairwise Wilcoxon test with Bonferroni correction); ns, not significant. ♢ Average per group **(d)** Scanning electron microscopy (SEM) of a germinated zoospore (g.z.) directly penetrating the root (white arrowhead) at 1dpi, scale bar: 10 μm **(e)** SEM of root surface colonized by hyphae at 3 dpi, scale bar: 200 µm. **(f)** SEM of *A. euteiches* oospores (white arrowheads) on the surface of n°76 roots at 6 dpi, scale bar: 50 μm. **(g)** Cross-section of inoculated roots showing *A. euteiches* oospores (white arrowheads) at 6 dpi (brightfield microscopy), scale bar: 200 μm.

At 21 days post inoculation (dpi), inoculated n°76 plants displayed visible differences in root morphology compared to non-inoculated controls. The RB84 strain induced extensive root browning, a marked reduction in both primary root length and lateral root formation. In contrast, plants inoculated with the ATCC201684 strain exhibited milder symptoms, with less severe browning and relatively preserved root length. These visual observations suggested differences in virulence between the two strains (**Fig. 1a**).

To validate these observations, root browning and plant growth were quantified as described in the “Methods” section. The percentage of brown root surface was measured at 3, 5, 7, 10, and 12 dpi. Browning was observed as early as 3 dpi for both strains, indicating rapid pathogen colonisation and symptom development. However, plants inoculated with RB84 consistently exhibited a significantly higher percentage of browning than those inoculated with ATCC201684 at all time points (**Fig. 1b**; p <0.01 or < 0.001). In addition, pathogen impact on plant growth was quantified by measuring fresh weight at 21 dpi. A significant reduction in fresh weight was observed in plants inoculated with RB84 compared to non-inoculated controls (**Fig. 1c**; p < 0.001), whereas no significant difference was detected for plants inoculated with ATCC201684. This quantitative dataset confirms the higher virulence of RB84. Notably, no plant mortality was observed in any replicate, regardless of the strain, highlighting the partial resistance of *A. evenia* to *A. euteiches*.

To further assess the compatibility of the interaction, root samples were examined using scanning electron microscopy (SEM) (**Fig. 1d-f**). At 1 dpi, encysted zoospores had germinated and directly penetrated root epidermal cells (**Fig. 1d**) without formation of penetration structures. By 3 dpi, the root surface was covered with a hyphal network, with some hyphae emerging from the root cortex (**Fig. 1e**). Root hairs collapsed beneath the mycelium, and macerated areas were observed on the root surface, coinciding with the onset of browning (**Fig. 1e**). For both strains, SEM at 6 dpi and bright-field microscopy of cross-sections at 6 dpi revealed the presence of oospores on the root surface (**Fig. 1f**) and within the cortical tissue (**Fig. 1g**). At this stage, infection induced epidermal cell browning and the accumulation of brown cytoplasmic compounds in some cortical cells. These observations confirm successful infection and demonstrate that the pathogen completes its reproductive cycle on *A. evenia*.

Altogether, these findings demonstrate that the interaction between *A. evenia* and *A. euteiches* is compatible and that RB84 is more virulent than ATCC201684 on the n°76 genotype. The RB84 strain was therefore selected for subsequent experiments.

### 3.2 *A. euteiches* inoculation induced a transcriptomic response in *A. evenia*

Based on microscopic and phenotypic analyses, two time-points were selected for RNA-Seq analyses in the n°76 genotype to capture the dynamics of the early stages of the interaction: 1 dpi, representing the plant’s response to pathogen penetration (**Fig. 1d**), and 3 dpi, when cortical colonisation and root browning become visible. RNA from roots collected at T0 was used as a non-inoculated reference control. For each time-point four biological replicates of inoculated or mock-inoculated samples were analysed, generating on average 20.9 to 32.6 million reads per sample. Quality control of the normalised data, using multidimensional scaling (MDS) showed that non-inoculated and mock-inoculated samples clustered closely, whereas infected samples were clearly separated, particularly at 3 dpi, confirming infection-triggered transcriptomic changes (**Fig. 2a**). The impact of infection was stronger at 3 dpi than at 1 dpi and slight dispersion among replicates likely reflected differences in zoospore inoculum preparation.

**Figure 2:**
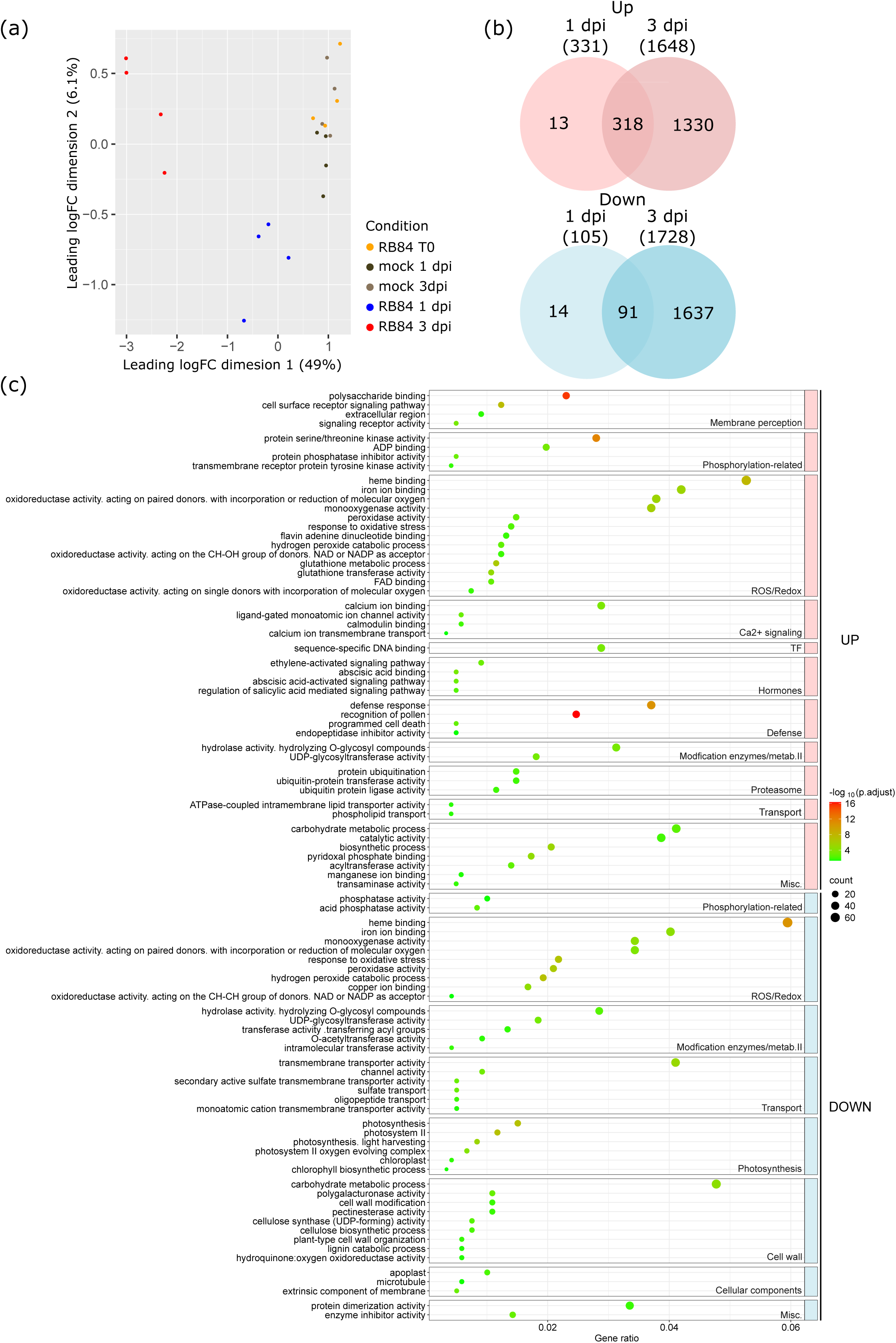
*A. euteiches* inoculation induces a transcriptomic response in *A. evenia*. **(a)** Multidimensional scaling (MDS) plot showing clustering of non-inoculated (T0) and mock-inoculated samples (1 dpi) on the left, and inoculated samples (1 dpi and 3 dpi) on the right. The plot illustrates transcriptomic separation between conditions, with dimension 1 explaining 49% of the variance and dimension 2 explaining 6.3%. **(b)** Numbers of differentially expressed genes (DEGs) in *A. evenia* roots at 1 and 3 days post-inoculation (dpi) with *A. euteiches*. DEGs were identified using a fold-change threshold of |logFC| ≥ 1 and a false discovery rate (FDR) < 0.05. Venn diagrams indicate the overlap and specificity of upregulated and downregulated genes at each time point. **(c)** Gene ontology (GO) enrichment analysis among all DEGs at 3dpi, separated into upregulated and downregulated groups. GOs are shown with their corresponding adjusted p-values and gene ratios, highlighting categories associated with immune signaling, metabolism, and transcriptional regulation.

RNA-Seq revealed significant transcriptomic reprogramming, with a total of 3,403 differentially expressed genes (DEGs) detected across the two time-points. At 1 dpi, 436 DEGs were identified (331 upregulated, 105 downregulated), whereas at 3 dpi, this number rose dramatically to 3,376 DEGs (1,648 upregulated, 1,728 downregulated) (**Fig. 2b and Supp data; Table S4**). Among the 1,661 upregulated genes, 318 were shared between time points, whereas only 91 downregulated genes overlapped. Most DEGs (87%) were specific to 3 dpi, reflecting a progressive and amplified response to the pathogen, with a majority (55%) being downregulated (**Fig. 2b**).

#### *A. evenia* activates an early immune response to *A. euteiches* infection

To dissect the transcriptional dynamics underlying the response of *A. evenia* to *A. euteiches*, we performed GO enrichment and InterPro domain analyses on DEGs at 1 dpi and 3 dpi **(Supp data; Table S7 and S8)**.

At 1 dpi, no GO term enrichment was detected among downregulated genes, suggesting an early reprogramming dominated by activation rather than repression. Functional categories highlighted pathogen perception and early immune signaling, including receptor-mediated recognition and signaling cascades, supported by the overrepresentation of leucine-rich repeat (LRR) domains, typically found in receptors recognising pathogen-derived peptides, as well as bulb-type lectins and Gnk2-like proteins known to bind pathogen-associated carbohydrates (Miyakawa *et al*., 2014; Zou *et al*., 2022). The enrichment of kinase and wall-associated kinase domains further reflects the activation of intracellular signaling through phosphorylation.

In parallel, genes involved in redox regulation were induced, with oxidoreductases, glutathione-related enzymes and thiolases suggesting an early adjustment of ROS homeostasis. This response was accompanied by the induction of transcriptional regulators such as WRKY and ethylene-responsive transcription factors, consistent with their known roles in immune gene expression (Birkenbihl *et al*., 2017; Dubois *et al*., 2018). Hormonal signaling components related to abscisic acid and ethylene were also detected.

Specialised metabolism was activated early, with induction of several enriched domains, including chalcone/stilbene synthase, cytochrome P450 or polyketide synthase, consistent with the biosynthesis of antimicrobial compounds. Finally, the enrichment of transporter-related GO terms, together with domains linked to amino acid and heavy metal transporters, suggests adjustments in nutrient fluxes upon infection.

Altogether, these enrichments reflect a multifaceted early immune response at 1 dpi, which later expands into a broader and more integrated immune program at 3 dpi.

#### *A. evenia* exhibits an expanded immune and metabolic response at 3 dpi

At 3 dpi, the number of DEGs increased dramatically compared to 1 dpi, and the enriched functional categories pointed to both an amplification and a diversification of immune responses (**Fig. 2b-c and Supp data; Table S7**). Categories associated with early perception and membrane signaling, including receptor activities and phosphorylation events, were still detected, indicating sustained surveillance as the pathogen entered the root cortex. In addition, enriched IPR domains such as NB-ARC domains and Toll/interleukin receptor domains (**Supp data; Table S8**) were overrepresented. The detection of these resistance gene hallmarks (Castel *et al*, 2024), suggests that, beyond the PRR-mediated perception observed at 1 dpi, *A. evenia* mobilizes intracellular NLR receptors at 3 dpi to reinforce effector recognition.

A prominent feature of the 3 dpi response was the regulation of oxidative stress. More than 12 GO terms, including response to oxidative stress, hydrogen peroxide catabolism, heme binding, peroxidase activity and glutathione metabolism, were significantly enriched among both up- and downregulated genes. Consistently, domains related to ROS detoxification, such as haem peroxidase, multicopper oxidases, dioxygenases, and NAD(P)-binding domains were overrepresented (**Supp data; Table S8**). This points to an intensified regulation of ROS homeostasis, balancing their dual role as signaling molecules and cytotoxic agents, and requiring detoxification systems from the plant (Waszczak *et al*., 2018). Calcium-related categories also emerged, consistent with the enrichment of EF-hand domains, typical of calcium sensors such as calmodulin. In parallel, signaling linked to Abscisic acid (ABA) and ethylene (ET) pathways was maintained and complemented by induction of the salicylic acid (SA) pathway, reflecting an amplification of immune signaling.

Beyond perception and signaling, downstream defence mechanisms were also activated. Functional enrichments included PR-proteins, such as chitinases, β-1,3-glucanases, and thaumatin-like proteins, all contributing to pathogen cell wall degradation. In parallel, enzymes of secondary metabolism - including chalcone/stilbene synthases, polyketide synthases, and cytochrome P450 monooxygenases - were induced, consistent with flavonoid and phytoalexin biosynthesis. Together, these enrichments indicate the deployment of an antimicrobial machinery, combining PR-protein activity with the production of specialised metabolites, thereby reinforcing the chemical barriers deployed by *A. evenia* against *A. euteiches*.

Beyond canonical immunity, 3 dpi was also marked by extensive metabolic adjustments. Categories included transferases, hydrolases, dioxygenases and carbohydrate metabolism, supported by the induction of glycosidase and α/β hydrolase domains. The activation of proteasome-related functions, together with the regulation of multiple transporters, underscores the broad cellular dynamics associated with infection and defence-related remodelling.

#### Growth repression and metabolic trade-offs

In contrast to 1 dpi, repression was markedly more pronounced at 3 dpi, with numerous enriched categories linked to growth and primary metabolism. Downregulated functions included cell wall organization or pectin catabolism, reflected by decreased representation of pectin lyases, arabinogalactan proteins, fasciclin-like domains, and pectin esterase inhibitors. Several categories related to sugar metabolism, glycosyltransferases or cellulose synthases were also repressed. A consistent trend was the downregulation of photosynthesis-related functions, including chlorophyll a/b-binding proteins, as well as reductions in domains associated with chloroplast functions and the cytoskeleton (**Fig. 2c and Supp data; Table S8**) pointing to a strong trade-off between growth and defence (Berens *et al*., 2017).

To conclude, infection of the *A. evenia* n°76 genotype by *A. euteiches* at 3 dpi triggered transcriptomic changes consistent with canonical immune responses, including genes implicated in pathogen perception, signaling, transcriptional regulation, defence, metabolic reprogramming, cell wall remodelling and oxidative stress management. Altogether, enrichments at 3 dpi reveal that the initial immune responses observed at 1 dpi are not only maintained but are rapidly complemented by new layers of defence, including NLR activation, enhanced regulation of ROS homeostasis, intensified Ca²⁺ and hormone signalling, large-scale metabolic reprogramming, and repression of photosynthesis and growth-related functions. These results highlight an amplified and integrated plant response to counteract pathogen invasion within the root cortex.

### 3.3 Comparative transcriptomic responses of Aeschynomene evenia and Medicago truncatula to Aphanomyces euteiches

To conduct a comparative transcriptomic analysis, the set of DEGs identified in *A. evenia* n°76 at 3 dpi with *A. euteiches* was compared to the 2,547 significant DEGs obtained from an RNA-Seq experiment on root tissues of *M. truncatula* line A17 inoculated with *A. euteiches*. This *Medicago* line has previously been described as partially resistant and has been used to investigate mechanisms underlying resistance to Aphanomyces root rot (ARR) (Djébali *et al*., 2009; Rey *et al*., 2013; Badis *et al*., 2015). Root tissues from both species were harvested at 3 dpi after inoculation with the virulent *A. euteiches* strain RB84. Among upregulated genes, 2,239 were identified in *M. truncatula*, and 1,648 in *A. evenia,* indicating roughly similar orders of magnitude (**Fig. 3a**). By contrast, only 308 downregulated genes were detected in *M. truncatula*, compared to 1,728 in *A. evenia*, revealing a major quantitative difference between the two species (**Fig. 3a**).

**Figure 3:**
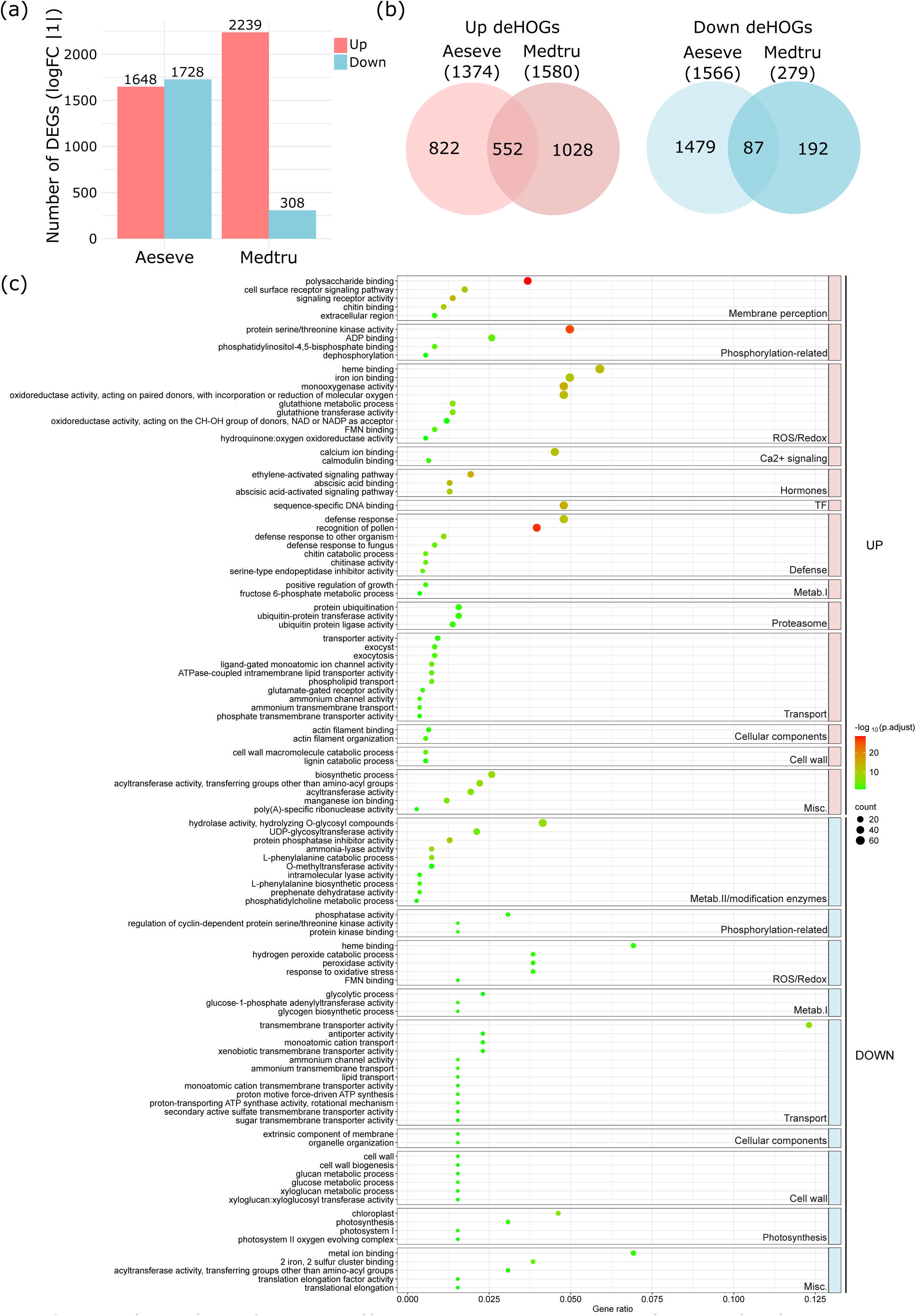
*A. euteiches* inoculation induces a comparable transcriptomic response in *A. evenia* and *M. truncatula*, with quantitative differences in gene repression. **(a)** Number of differentially expressed genes (DEGs) in *A. evenia* and *M. truncatula* at 3 dpi. DEGs were identified using a fold-change threshold of |logFC| > 1and a false discovery rate (FDR) < 0.05. **(b)** Numbers of hierarchical orthogroups (HOGs) containing DEGs in *A. evenia* and *M. truncatula* roots at 3 days post-inoculation (dpi) with *A. euteiches*. DEGs were identified using a fold change threshold of |logFC| > 1 and a false discovery rate (FDR) < 0.05. Only HOGs in which DEGs were homogeneously regulated (either up- or downregulated) within the orthogroups were considered. Venn diagrams indicate the overlap of such differentially expressed HOGs (deHOGs) between the two species. **(c)** Gene ontology (GO) term enrichment among common dysregulated HOGs separated into upregulated and downregulated groups. GOs are represented with their corresponding adjusted p-values and gene ratios.

To allow cross-species comparisons, DEGs were then assigned to hierarchical orthologous groups (HOGs), as described in the Methods and Supplementary Methods sections. Comparisons of up- and downregulated HOGs in both species are presented in **Fig. 3b**. This analysis revealed a substantial overlap for upregulated HOGs (552 shared at 3 dpi), whereas only a limited number (87) were shared among the downregulated HOGs (**Fig. 3b**). To further characterise the conserved response between the two species, we analysed GO enrichment among the shared up- and downregulated HOGs.

Enrichment among the shared upregulated HOGs highlighted a broad activation of immune-related processes, including pathogen perception, early signalling (kinase activity, calmodulin binding, ET- and ABA-related pathways), and redox homeostasis (oxidoreductases, heme binding, monooxygenases) (**Fig. 3c**). Additional enrichments pointed to ubiquitin-mediated protein degradation, carbohydrate metabolism (glycoside hydrolases, glutathione transferases), and the biosynthesis of specialised metabolites, consistent with a conserved immune activation signature in both species.

In contrast, the set of commonly downregulated HOGs was smaller but pointed to similar physiological shifts. Suppressed categories included redox- and oxygen-related metabolism (peroxidases, heme binding), diverse transport functions (sugar, cation and ammonium transporters), photosynthesis-related functions (chloroplast proteins, photosystem II components), and cell wall modification (glucan and xyloglucan metabolism, transferases).

Together, these patterns indicate that, despite quantitative differences, *A. evenia* and *M. truncatula* undergo qualitatively similar transcriptional reprogramming, characterised by the induction of defence-related functions and trade-offs in carbon metabolism, cellular dynamics, and cell-wall remodelling.

### 3.4 Diversity of quantitative disease resistance levels among natural accessions of *A. evenia*

To explore natural variation in plant responses to *A. euteiches*, five *A. evenia* accessions (n°76; n°77; n°79; n°80; n°21) were inoculated with the RB84 strain. Symptoms were monitored at multiple time points (3, 5, 7, 10, and 12 dpi), and the area under the curve (AUC) of the percentage of root-surface browning was calculated to integrate the kinetics of symptom development (**Fig. 4a**). This analysis revealed significant variability in resistance levels among accessions. Accession n°76 showed the highest levels of root browning and was the most susceptible, whereas n°79 displayed the least browning and was identified as the most resistant. Accessions n°21, n°77, and n°80 displayed intermediate phenotypes.

**Figure 4:**
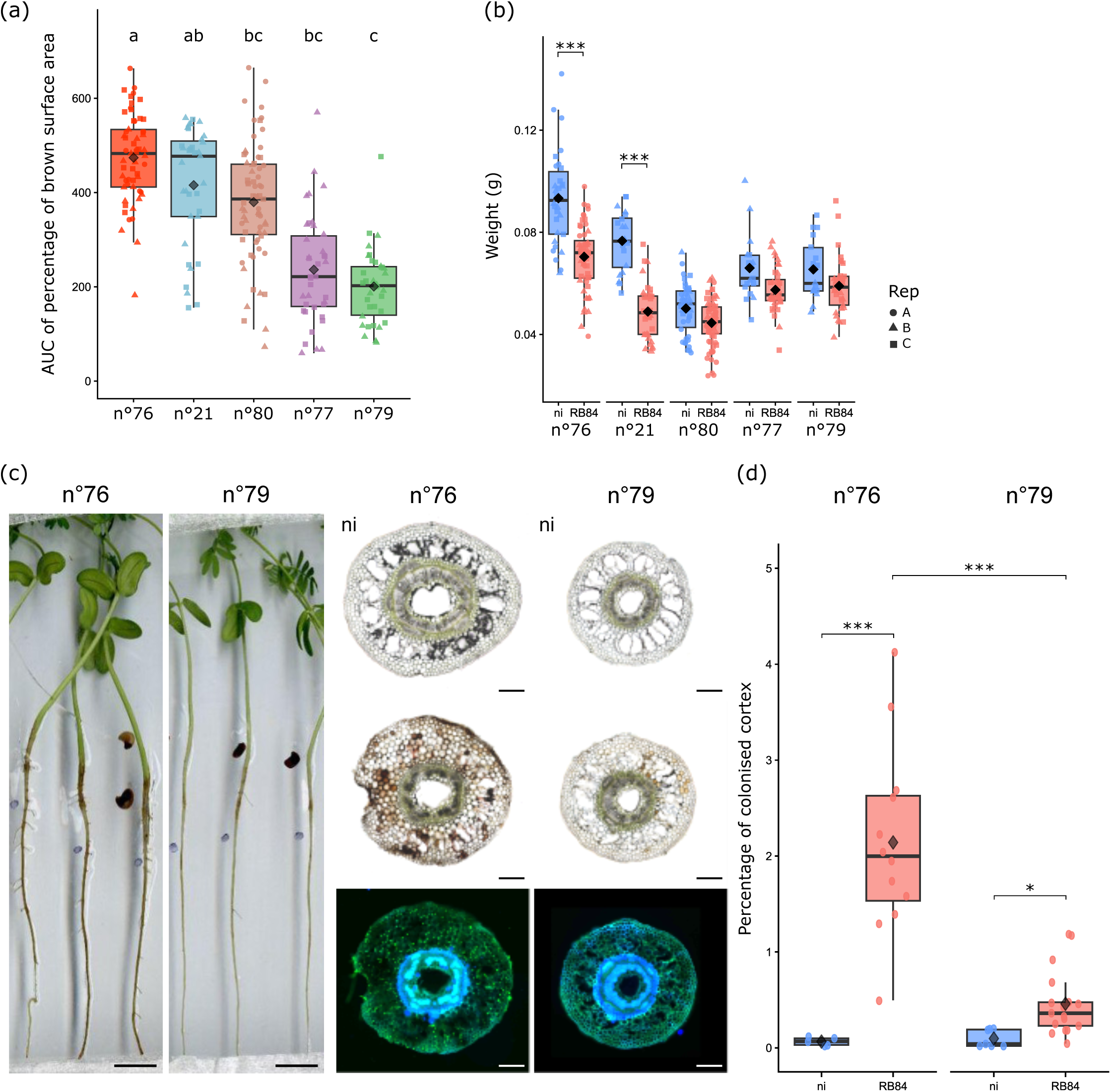
Natural diversity of quantitative disease resistance to *A. euteiches* across *A. evenia* accessions. **(a)** Area under the curve (AUC) of percentage of root surface browning measured over a time course of infection at 3, 6, 9, and 12 dpi. Statistical differences between accessions were determined using pairwise Wilcoxon tests with Bonferroni correction (a-b: p = 0.03; a-c: p = 0.0021); N = 41-45 plants, depending on the *A. evenia* accession. ♢ Average per group **(b)** Fresh weight of non-inoculated (ni) and inoculated plants at 21 days post-inoculation (dpi) with the RB84 strain. Statistical differences were analyzed using pairwise Wilcoxon tests with Bonferroni correction (***p < 0.001). N = 12-15 plants for non-inoculated controls; N = 26-30 for inoculated plants. ♢ Average per group **(c)** Phenotypes and cytological analyses of n°76 and n°79 *A. evenia* accessions after A. euteiches infection. Left panel: n°76 & n°79 root scans at 6 dpi show stronger and faster development of root browning in n°76 compared to n°79. Right panel: Thin root sections (100 µm) of non-inoculated (ni) and inoculated roots with the RB84 strain at 21 dpi, observed under fluorescence microscopy. A higher hyphal density is observed in the cortical cells of n°76 compared to n°79. *A. euteiches* mycelium is labeled in green using WGA-FITC staining. Scale bar = 100 µm. Brightness increased by 10% in BF, 50% in fluorescence, contrast by 20% in fluorescence. **(d)** Percentage of colonised cortex surface, calculated as the ratio of WGA-FITC fluorescent area to the total cortex area. Statistical differences were analyzed using pairwise Wilcoxon tests with Bonferroni correction (***p < 0.001). N = 12-15 sections for non-inoculated controls; N =26-30 section for inoculated plants. ♢ Average per group

The resistance gradient was confirmed by measuring plant fresh weight at 21 dpi (**Fig. 4b**). Compared to non-inoculated controls, inoculated plants of n°76 and n°21 showed significant reductions in fresh weight, while n°77, n°80, and n°79 displayed only minor, non-significant decreases (**Fig. 4b**). No mortality was observed in any accession, underscoring that resistance levels varied quantitatively rather than qualitatively.

Two contrasting accessions, n°76 (susceptible) and n°79 (resistant), were selected for cytological analyses. At 6 dpi, root scans showed earlier and more extensive browning in n°76 compared to n°79 (**Fig. 4c**). At 21 dpi, fluorescence microscopy revealed intracellular colonisation of cortical tissues by *A. euteiches* in both genotypes, but hyphal density was much higher in n°76. Quantification of colonised cortical surface confirmed a significantly lower infection level in n°79 compared to n°76 (**Fig. 4d**). In addition, increased autofluorescence of endodermal cell walls and pericycle cell divisions were observed, indicating cellular responses to pathogen invasion.

Altogether, these results underscore the diversity of resistance levels among *A. evenia* accessions, evident both at the macroscopic level and in their cellular responses to *A. euteiches* colonisation.

### 3.5 An overlapping response between infection with *A. euteiches* and NF-independent symbiosis with *Bradyrhizobium*

To leverage this newly established *A. evenia–A. euteiches* pathosystem, we examined whether immunity and NF-independent symbiosis share common transcriptional features. To do so, we compared the *A. evenia* transcriptomic response to *A. euteiches* with its response to the Nod factor-independent symbiont *Bradyrhizobium* ORS278, described by Gully *et al*. (2018). Across the different time points after inoculation with *Bradyrhizobium spp.*, 3623 genes were identified as significantly differentially expressed. Comparing these DEGs with the 3403 DEGs detected in response to *A. euteiches* inoculation identified a core set of 1,103 DEGs shared across both interactions, including 215 commonly upregulated and 517 commonly downregulated genes (**Fig. 5a**). A heatmap of all genes significantly differentially expressed in either response reveals contrasting expression profiles, especially between DEGs induced by *A. euteiches* at 3dpi and those responding to *Bradyrhizobium spp.* at 144 hpi (**Fig. 5b**). Clustering of conditions revealed that the most similar transcriptomic responses to both microorganisms occurred at 1dpi, after which the two interactions diverged markedly over time. Contrasted profiles consisted of 164 genes upregulated in response to *Bradyrhizobium spp.* and downregulated in response to *A. euteiches,* and 207 genes showing the opposite pattern.

**Figure 5:**
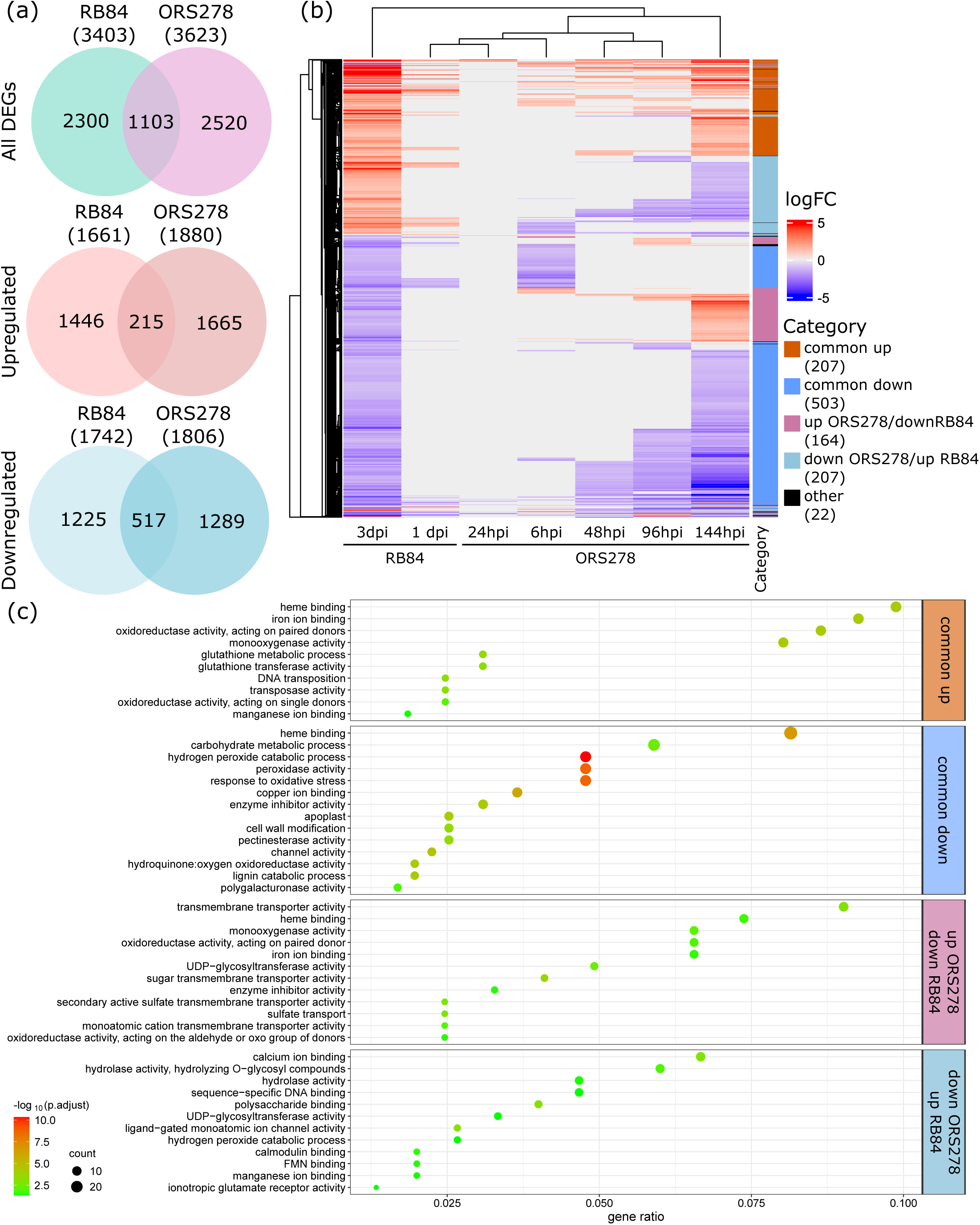
Common and specific genes are differentially expressed in response to *A. euteiches* inoculation and NF-independent symbiosis. **(a)** Numbers of differentially expressed genes (DEGs) in *A. evenia* roots post-inoculation (dpi) with the RB84 *A. euteiches* strain or the *Bradyrhizobium* ORS278 strain. DEGs were identified using a fold-change threshold of |logFC| > 1 and a false discovery rate (FDR) < 0.05. Venn diagrams indicate the overlap and specificity of deregulated, upregulated and downregulated genes for each inoculated microbe. **(b)** Clustered heatmap of gene significantly differentially expressed in response to *Bradyrhizobium spp.* (ORS278) and *A. euteiches* (RB84) accompanied with their corresponding LogFC for each timepoint and condition. **(c)** Significantly enriched gene ontology (GO) terms among common and opposite profile DEGs. GO terms are shown with their corresponding adjusted p-values and gene ratios, highlighting functional categories associated mainly with ROS homeostasis and cell wall remodeling.

#### Shared responses

GO enrichment among the shared upregulated genes (**Fig. 5c**) highlighted a strong induction of ROS and redox-related functions, including oxidoreductase activity, heme and iron ion binding, monooxygenase activity, and glutathione metabolism. Consistently, the InterPro analysis showed enrichment of domains directly associated with ROS regulation and detoxification, such as thioredoxin-like, and glutathione S-transferase (**Supp data; Table S10)**. Additional enriched functions included DNA transposition, suggesting a potential involvement of transposable elements in transcriptional plasticity. Furthermore, Interpro domains linked to microbial perception and specialised metabolism (e.g. cytochrome P450s, aminotransferases) were also enriched, indicating that the common response combines redox regulation with the mobilisation of metabolic pathways involved in biotic interactions.

Conversely, GO terms enriched among the shared downregulated genes indicated broad suppression of growth-, metabolic-, and transport-related processes. This included carbohydrate metabolism and cell wall remodelling, reflected by enrichments in pectinesterases, polygalacturonases, and related domains. A second group of enriched terms related to ROS and redox processes - such as peroxidase activity, hydrogen peroxide catabolism, and copper ion binding was consistent with the Interpro enrichment of multicopper oxidases (cupredoxin and secretory peroxidase domains) (**Supp data; Table S10)**. Finally, transport-related functions, including channel activity, were also repressed, matching decreases in aquaporin- and oligosaccharide transporter-associated domains.

Together, these results highlight the central role of ROS homeostasis in both pathogenic and symbiotic interactions, involving coordinated up- and downregulation of redox-associated functions. In addition, the shared transcriptional signature points to convergent changes in specialised metabolism, transport, and cell-wall remodelling - processes likely contributing to the regulation of microbial progression within root tissues.

#### Contrasted responses

Among contrasted regulation profiles, genes positively regulated in response to *Bradyrhizobium spp.* present an enrichment of terms associated with transmembrane transport and sugar and sulfate transporters along with REDOX regulation (**Fig 5c**). The importance of transporters among up regulated genes in symbiosis is reinforced in the Interpro domain enrichment, containing SWEET domains along with MFS and other transporter families (**Table S10)**.

On the other hand, genes upregulated in response to *A. euteiches* but repressed during symbiosis displayed a transcriptional signature fully consistent with immune activation. GO enrichment highlighted strong induction of calcium- and calmodulin-binding activities, ROS-associated processes such as hydrogen-peroxide catabolism and oxidoreductase activity, and DNA-binding functions linked to transcriptional reprogramming.

Consistently, InterPro enrichment revealed domains encoding key components of these immune modules, including EF-hand and calmodulin-like domains involved in Ca²⁺ sensing, germin/cupin and peroxidase families associated with ROS production and detoxification, nitric-oxide-synthase-like signatures, and transcription-factor domains such as WRKY.

Together, these features show that genes activated during immunity but suppressed during symbiosis engage interconnected Ca²⁺-dependent signalling, redox regulation and transcription-factor activity—three hallmarks of canonical defence activation in plants.

Overall, the enriched GO and InterPro terms among genes with contrasted expression profiles highlight a focus on transport and exchange during symbiosis and on perception and immune signalling in immunity, both interactions are underpinned by pervasive regulation of redox homeostasis.

### 3.6 *AeRLCK2* is a negative regulator of resistance to *A. euteiches*

Among the genes responding to *A. euteiches*, kinases were particularly enriched in the*A. evenia* transcriptome, suggesting a central role for early signalling components in the interaction. We therefore focused on *AeRLCK2* (*Ae01g26600*), a receptor like cytoplasmic kinase required for nodule initiation in the Nod factor-independent symbiosis and its known interactor *AeCRK* (*Ae05g12380*), another RLK (Quilbé *et al.,* 2021, Horta Araújo *et al*., 2025). Transcriptomic data indicated that *AeRLCK2* was significantly induced at 3 dpi with *A. euteiches* (logFC ≈ 0.96, p ≈ 5 × 10⁻⁶, whereas *AeCRK* did not show significant differential expression - (**Table S4**), To determine whether these kinases contribute to the plant response to *A. euteiches*, EMS-induced allelic mutants of *AeRLCK2* (*rlck2-1* and *rlck2-2*) and *AeCRK* (*crk-*1 and *crk-*2) were inoculated with *A. euteiches* and compared to the wild-type (WT) reference line n°76.

Analysis of the area under the curve (AUC) of percentage of root-surface browning (**Fig. 6a**) revealed that both *AeRLCK2* and *AeCRK* mutants exhibited significantly reduced browning compared with WT. Fresh weight measurements at 21 dpi showed a significant reduction upon infection, in the WT and in the *crk-2* mutant whereas *rlck2-1*, *rlck2-2* and *crk-1* displayed only minor, non-significant decreases, thereby supporting deeper focus on the *rlck2* mutants (**Fig. 6b**). At the microscopic level, image-based quantification of hyphal colonisation (**Fig. 6c**) revealed a lower percentage of colonised cortical tissue in *rlck2* mutant root sections than in WT (n°76) (**Fig. 6d**).

**Figure 6:**
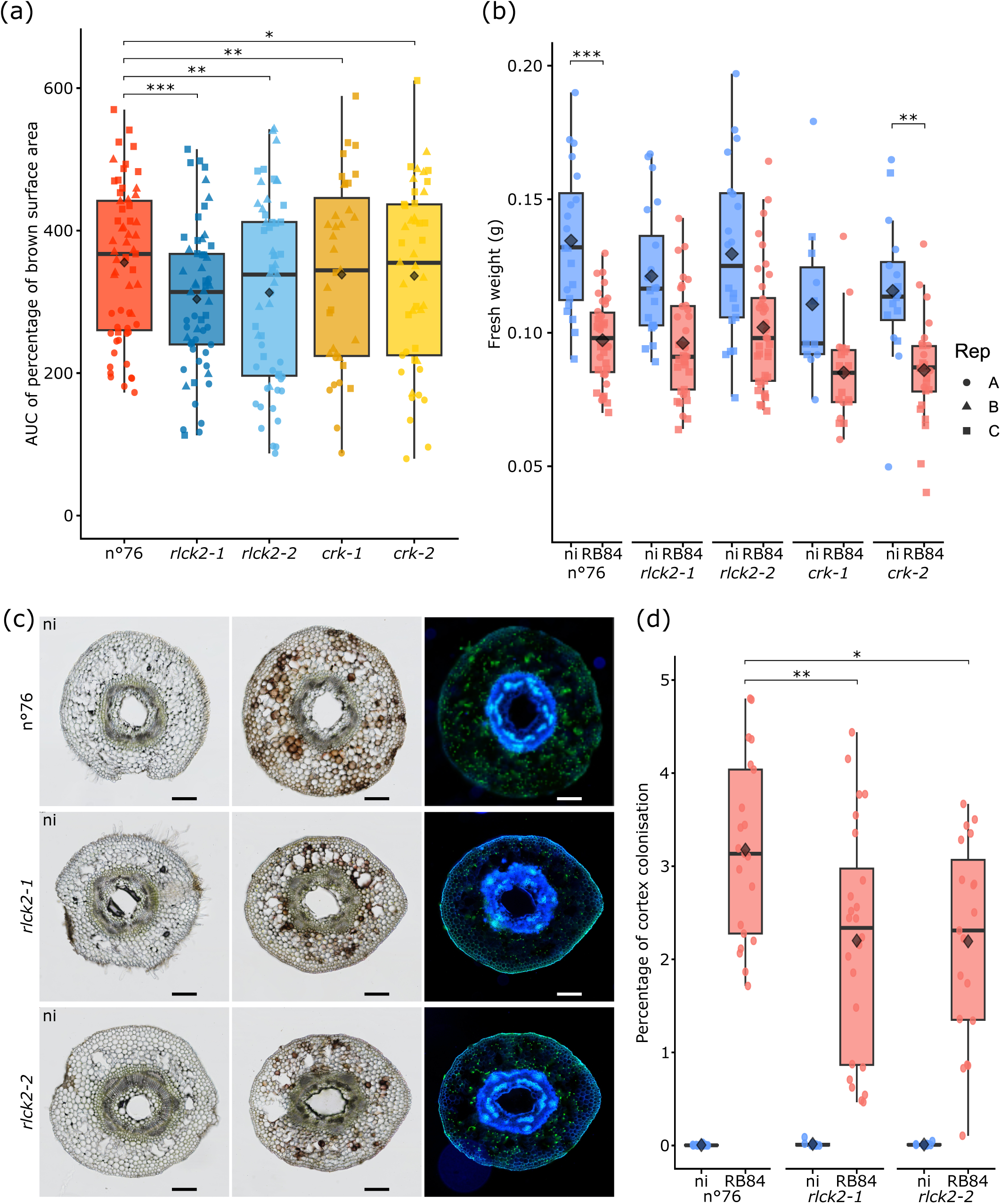
Nod Factor-independent symbiotic mutants exhibit increased resistance to *A. euteiches*. **(a)** Area under the curve (AUC) of percentage of root surface browning measured over a time course of infection at 3, 5, 7, 10, and 12 dpi. Kinase mutants in the Nod Factor-independent symbiotic pathway exhibit varying levels of susceptibility to *A. euteiches*. Statistical differences were analyzed using a linear model to compare means of all allelic mutants to the WT (n°76) considering replicate effect (*p < 0.05, **p < 0.01, ***p < 0.001) N = 45 plants per accession. **(b)** Fresh weight of non-inoculated and inoculated plants at 21 dpi. Symbiotic mutants exhibited different levels of growth reduction in response to *A. euteiches* infection. Statistical differences were analyzed using pairwise Wilcoxon tests with Bonferroni correction (*p < 0.05, **p < 0.01, *** p<0.001). **(c)** Cytological analysis of mutant colonisation. Thin root sections (100 µm) of non-inoculated (ni) and inoculated roots with the RB84 strain at 21 dpi, observed under fluorescence microscopy. A higher hyphal density is observed in the cortical cells of n°76 compared to rlck2 mutants. *A. euteiches* mycelium is labeled in green using WGA-FITC staining. Scale bar = 100 µm. **(d)** Percentage of colonised cortex surface of non-inoculated and inoculated plants at 21 dpi, calculated as the ratio of WGA-FITC fluorescent area to the total cortex area. Statistical differences were analyzed for each mutant line using Wilcoxon tests with Bonferroni correction (*p < 0.05, **p < 0.01).

Together, these macro- and microscopic analyses, combined with the induction of *AeRLCK2* during infection, suggest that this kinase contributes to the modulation of quantitative resistance to *A. euteiches*. The enhanced resistance observed in the knockout rlck2 mutants further indicates that AeRLCK2 acts as a negative regulator of immunity, or alternatively, that it is required for efficient colonisation of root tissues by *the oomycete*. *Aecrk* mutants displayed a similar, but less pronounced phenotype. This result may reflect either a milder loss of function in the available alleles or a less central role of *AeCRK* in modulating resistance compared with *AeRLCK2*.

## 4 Discussion

This work establishes *Aeschynomene evenia* as a novel pathosystem to investigate quantitative disease resistance (QDR) against the soilborne oomycete *Aphanomyces euteiches.* By combining phenotypic, cytological and transcriptomic analyses with comparative approaches, and functional analyses with symbiotic mutants, our study provides insights into the dynamics of immune responses and their potential overlap with Nod factor-independent symbiosis (NFIS). The implications of these findings are considered in light of previous work on legumes and plant–oomycete interactions, emphasizing both conserved and specific aspects of *A. evenia* immunity, including the emerging dual role of AeRLCK2 and AeCRK.

### 4.1 The *A. evenia–A. euteiches* pathosystem: ecological relevance and potential as a model

*Aphanomyces euteiches* is a specialized pathogen of the Fabaceae family, affecting multiple crop and forage legumes such as pea, alfalfa, beans, lentils, clovers, vetches, and faba beans, predominantly in temperate regions of the Northern Hemisphere (Gaulin *et al*., 2007, Moussart *et al*., 2008; Becking *et al*., 2022). Our study demonstrates the compatibility between *A. euteiches* and *A. evenia*, with two strains of the pathogen completing their reproductive cycle, while resistance levels varied quantitatively across accessions. These findings establish *A. evenia* as a potential host for this oomycete, and the absence of full resistance combined with the lack of mortality is a hallmark of QDR, a prevalent form of resistance in legumes and other crops (Bonhomme *et al*., 2019; Roux *et al*, 2014).

However, *A. evenia* is a tropical or subtropical species with a semi-aquatic lifestyle. The natural accessions tested in this study originate from Southeast Africa (n°76 and n°21 lines) and South America (n°77, n°79, and n°80 lines) (Quilbé *et al*., 2021). Given the environmental preferences of *A. evenia* and the predominantly temperate distribution of *A. euteiches* (Becking *et al*., 2022), it is unlikely that these species naturally cohabit. Nevertheless, the humid and warm conditions under which *A. euteiches* thrives in our experiments (25°C in this study, but it can also grow without difficulty at 28°C, not shown) align with the ecological preferences of *A. evenia*, suggesting that interactions with related oomycetes may occur in its native environment. Interestingly, the accessions displaying higher resistance to *A. euteiches* (n°79, n°77, n°80) originate from South America. Although further studies are needed to confirm this trend, these observations raise the possibility that South American accessions may have evolved resistance traits against pathogens with similar infection strategies. Testing additional accessions from this region, as well as challenging *A. evenia* with tropical pathogens—such as *Phytophthora palmivora* or even *P. nicotianae*, which cause root rot on legumes in South America (Cabral *et al*., 2024) could help assess this hypothesis, and further elucidate the genetic basis of resistance traits.

### 4.2 *A. evenia* shows common and specific QDR traits compared to other legumes in response to *A. euteiches*

The phenotypic responses of *A. evenia* to *A. euteiches* inoculation revealed both commonalities and unique features compared to those in *M. truncatula* and *P. sativum*. Root browning and stunted growth, hallmark traits of soilborne oomycete infection, were observed in all three species and are consistent with earlier studies in *M. truncatula* (Djébali *et al*., 2009; Djébali *et al*., 2011) and *P. sativum* (Lavaud *et al*., 2015). The dark brown pigmentation of resistant *A. evenia* accessions resembles that of the partially resistant *M. truncatula* accession A17, in response to *A. euteiches* (Djébali *et al*., 2009; Djébali *et al*., 2011, Bonhomme *et al*., 2014). Quantitative resistance was evident in *A. evenia*, with clear differences in the rate and extent of root browning and in fresh weight reduction across accessions, mirroring patterns previously described in both *M. truncatula* (Bonhomme *et al*., 2014; Bonhomme *et al*., 2019) and *P. sativum* (Lavaud *et al*., 2015). The two strains of *A. euteiches* tested, RB84 and ATCC201684, exhibited distinct virulence levels in *A. evenia*, as previously observed in other legumes. However, it is worth noting that none of the *A. evenia* accessions displayed plant mortality, with either strain, in contrast to the high mortality rates reported in some *M. truncatula* and *P. sativum* accessions under comparable conditions (Bonhomme *et al*., 2014; Lavaud *et al*., 2015; Bonhomme *et al*., 2019). A broader panel of *A. evenia* accessions will nevertheless be required to determine whether this trend holds across the species.

Cytological analyses also revealed parallels with pea and *Medicago*, including intracellular hyphal proliferation in cortical cells and completion of the pathogen’s reproductive cycle. However, unlike these species, *A. evenia* accessions consistently prevented hyphal entry into the central cylinder. This correlates with the enhanced accumulation of fluorescent phenolic defence compounds, observed both inside cytoplasm and as structural reinforcements in endodermal cell walls. Such responses likely involve lignin deposition, as suggested by our RNA-seq data and previously described in *M. truncatula* (Djébali *et al*., 2009).

The higher level of resistance observed in *A. evenia* may also be attributed to its distinctive morphological adaptations, likely linked to its semi-aquatic environment. The presence of aerenchyma in the cortical layers may reduce hyphal density by limiting available colonisation sites. In addition, the central cavity of *A. evenia* roots, absent in *M. truncatula* and *P. sativum*, may facilitate oxygen circulation under flooded conditions while also acting as a physical barrier, that restricts pathogen spread. More accessions of *A. evenia* need to be analysed before definitively assessing the contribution of these structural traits.

Transcriptomic analyses revealed that *A. evenia* shares with *M. truncatula* the ability to mount a canonical immune response, encompassing pathogen perception, early and late signalling, ROS and redox homeostasis, and defence activation, with extensive reliance on specialised metabolism pathways, including phenylpropanoid and flavonoid biosynthesis. The strong enrichment in both up- and downregulated genes associated with ROS production and detoxification in *A. evenia* reflects the delicate regulation of ROS homeostasis linked to their dual role as toxic molecules and signalling mediators, a pattern reminiscent of the oxidative bursts and ROS-scavenging activities described in *M. truncatula* (Djébali et al., 2011; Truong et al., 2025) and in other legumes (Johansson *et al.,* 2025).

All these transcriptomic responses align with microarray experiments (Rey *et al*., 2013; Badis *et al*., 2015) and RNA-seq data (this study) from the resistant *M. truncatula* line inoculated with *A. euteiches*. Although both *M. truncatula* and *A. evenia* exhibit progressive transcriptomic reprogramming after one day, a distinctive feature of *A. evenia* is the marked downregulation of genes at 3 dpi. The enriched functional categories among these repressed genes point to pronounced metabolic trade-offs, stronger than those observed in *M. truncatula* or *P. sativum* (Kälin et al., 2024).

### 4.3 Transcriptomic crosstalk between symbiosis and immunity

The comparative analysis of pathogenic and symbiotic transcriptomes in *A. evenia* revealed a substantial overlap, with hundreds of genes commonly regulated in both contexts, as well as genes with opposite regulatory responses to each microorganism. The similarity of the early responses in symbiosis and immunity suggests an overall shared initial phase of biotic perception that then diverged to respond accordingly to either symbiont or pathogen. This transition suggests that the plant recognises the nature of the “invader” enabling an appropriate response to either “friend or foe”, and proposing an initial immune response along with an important role for genes implicated in perception and signalling.

ROS emerged as one of the most enriched functional categories shared between *A. evenia* responses to *A. euteiches* infection and symbiosis with *Bradyrhizobium*. The induction of genes associated with both ROS production (oxidoreductases, peroxidases) and detoxification (glutathione metabolism, dioxygenases) reflects the fine balance required to harness their dual role as antimicrobial agents and signalling molecules. This duality is well established in plant–pathogen interactions (Waszczak *et al*., 2018), and our results in *A. evenia* fit within recent publications showing that ROS and associated redox pathways function as central regulators linking immune and symbiotic processes in legumes (Trương et al., 2025; Johansson *et al*., 2025).

Genes related to cell wall metabolism, including lignin biosynthesis, pectin modification, and structural glycoproteins, were also significantly enriched in the shared transcriptomic response. These data underline the pivotal role of the cell wall as a dynamic and remodelled interface during plant–microbe interactions. Depending on the context, the wall must either be strengthened through lignin deposition (Djébali *et al*., 2009; Djébali *et al*., 2011) or modulated at the level of pectin methylation, while in other cases it is weakened or remodelled by cell-wall-degrading enzymes to facilitate microbial entry (Zhao *et al*., 2025; Tsyganova *et al*., 2021). In legumes, such remodelling is central to symbiotic infection modes: whereas most species rely on infection threads, *A. evenia* establishes Nod factor-independent symbiosis through *crack entry* at lateral root bases. This process also involves localised loosening and restructuring that allow bacterial progression (De Carvalho-Niebel *et al*., 2024). The convergence of cell-wall-related functions in both pathogenic and symbiotic contexts therefore suggests that *A. evenia* deploys overlapping regulatory modules to either restrict or accommodate microbial partners, depending on the interaction.

### 4.4 AeRLCK2 as a dual regulator of symbiosis and immunity in *A. evenia*

In *A. evenia*, AeRLCK2 has been recently identified as an essential component of Nod factor-independent symbiosis - NFIS - (Horta Araújo *et al*., 2025). Although its interactor AeCRK also contributes to NFIS, the stronger resistance observed in *Aerlck2* mutants, along with the significant induction of *AeRLCK2* during *A. euteiches* infection, suggest that AeRLCK2 plays the predominant role in this pathosystem. This dual implication (symbiosis establishment and decrease of plant immunity) is different from that of other RLKs, shown to be positive regulators of both symbiosis and immunity (Rey *et al*., 2013; Gibelin-Viala *et al*., 2019; Sarrette *et al*., 2025). It is, however, somehow reminiscent of the symbiotic receptor SymRK, which attenuates PTI through its interaction with LjBAK1 to promote rhizobial infection in *Lotus japonicus* (Feng et al., 2021). By analogy, AeRLCK2 may act as a negative regulator of immune activation, fine-tuning defence intensity according to microbial context. This duality reinforces the emerging view that RLCKs act as central hubs where symbiotic and immune signalling converge, enabling plants to finely tune their responses to beneficial and pathogenic microbes (Wang *et al*., 2025).

AeRLCK2 may also contribute to the regulation of growth–defence trade-offs, a hallmark of RLCK function in other species. In rice, OsRLCK176 acts as a negative regulator of immunity whose stabilisation coordinates immune activation with developmental constraints (Mou et al., 2024). The extensive repression of growth-related pathways observed in *A. evenia* at 3 dpi suggests that similar regulatory principles may operate during oomycete infection. Although the underlying mechanisms remain unknown, AeRLCK2 emerges as a promising node for integrating immune signalling with the metabolic costs of defence.

### 4.5 Concluding remarks

Altogether, this study establishes *A. evenia–A. euteiches* interaction as a new and tractable legume pathosystem for exploring quantitative disease resistance. By combining high-resolution cytology, comparative transcriptomics and functional genetics, we identify AeRLCK2 and AeCRK as promising regulators operating at the crossroads of symbiosis and immunity. These findings open new avenues for dissecting the genetic and evolutionary bases of QDR in legumes and for understanding how NF-independent symbiosis interfaces with defence signalling.

## Acknowledgements

This work was funded by the Agence Nationale de la Recherche (ANR-DUALITY project, ANR-20-CE20-0017-01). This study is set in the framework of the “Laboratoire d’excellence (LABEX) TULIP (ANR 10 LABX 41) and of the Ecole Universitaire de Recherche (EUR) TULIP GS (ANR 18 EURE 0019). MBa acknowledges receipt of a PhD grant from the Ministère de l’Enseignement Supérieur et de la Recherche, France. We acknowledge the TRI-FRAIB imaging facility, member of the national infrastructure France-BioImaging supported by the French National Research Agency (ANR-10-INBS-04).

## References

Achari SR, Kaur JK, Mann RC, Sawbridge T, Summerell BA, Edwards J (2021) Investigating the effector suite profile of Australian *Fusarium oxysporum* isolates from agricultural and natural ecosystems. Plant Pathology 70, 387–396.

Andrews S (2010) FastQC: A Quality Control Tool for High Throughput Sequence Data. Available at: https://www.bioinformatics.babraham.ac.uk/projects/fastqc/

Arrighi JF, Cartieaux F, Brown SC, Rodier-Goud M, Boursot M, Fardoux J, Patrel D, Gully D, Fabre S, Chaintreuil C et al. (2012) *Aeschynomene evenia*, a model plant for studying the molecular genetics of the Nod-independent rhizobium–legume symbiosis. Molecular Plant-Microbe Interactions 25, 851–861.

Badis Y, Bonhomme M, Lafitte C, Huguet S, Balzergue S, Dumas B, Jacquet C (2015) Transcriptome analysis highlights preformed defences and signaling pathways controlled by the *prAe1* QTL, conferring partial resistance to *Aphanomyces euteiches* in *Medicago truncatula*. Molecular Plant Pathology 16, 973–986.

Bardou P, Mariette J, Escudié F, Djemiel C, Klopp C (2014) Jvenn: An interactive Venn diagram viewer. BMC Bioinformatics 15, 293.

Barrett T, Dowle M, Srinivasan A, Gorecki J, Chirico M, Hocking T, Schwendinger B, Krylov I (2025) data.table: Extension of “data.frame” (version 1.17.8). CRAN: Contributed Packages. Available at: https://CRAN.R-project.org/package=data.table

Bécard G, Fortin JA (1988) Early events of vesicular–arbuscular mycorrhiza formation on Ri T-DNA-transformed roots. New Phytologist 108, 211–218.

Becking T, Kiselev A, Rossi V, Street-Jones D, Grandjean F, Gaulin E (2022) Pathogenicity of animal and plant parasitic *Aphanomyces* spp. and their economic impact on aquaculture and agriculture. Fungal Biology Reviews 40, 1–18.

Berens ML, Berry HM, Mine A, Argueso CT, Tsuda K (2017) Evolution of hormone signaling networks in plant defence. Annual Review of Phytopathology 55, 401–425.

Birkenbihl RP, Liu S, Somssich IE (2017) Transcriptional events defining plant immune responses. Current Opinion in Plant Biology 38, 1–9.

Bonhomme M, André O, Badis Y, Ronfort J, Burgarella C, Chantret N, Prosperi JM, Briskine R, Mudge J, Debellé F et al. (2014) High-density genome-wide association mapping implicates an F-box encoding gene in *Medicago truncatula* resistance to *Aphanomyces euteiches*. New Phytologist 201, 1328–1342.

Bonhomme M, Fariello MI, Navier H, Hajri A, Badis Y, Miteul H, Samac DA, Dumas B, Baranger A, Jacquet C et al. (2019) A local score approach improves GWAS resolution and detects minor QTL: application to *Medicago truncatula* quantitative disease resistance to multiple *Aphanomyces euteiches* isolates. Heredity 123, 517–531.

Broad Institute (2019) Picard Toolkit. Available at: https://broadinstitute.github.io/picard/

Brottier L, Chaintreuil C, Simion P, Scornavacca C, Rivallan R, Mournet P, Moulin L, Lewis GP, Fardoux J, Brown SC et al. (2018) A phylogenetic framework of the legume genus *Aeschynomene* for comparative genetic analysis of Nod-dependent and Nod-independent symbioses. BMC Plant Biology 18, 44.

Cabral CS, Gandara ADCF, Ribeiro Martins FHS, Barboza EA, Rossato M, Reis A (2024) *Phytophthora* species causing root and crown rot on castor bean (*Ricinus communis*) in Brazil. Journal of Phytopathology 172, e13337.

Castel B, Mahboubi KE, Jacquet C, Delaux PM (2024) Immunobiodiversity: conserved and specific immunity across land plants and beyond. Molecular Plant 17, 92–111.

Chaintreuil C, Perrier X, Martin G, Fardoux J, Lewis GP, Brottier L, Rivallan R, Gomez-Pacheco M, Bourges M, Lamy L et al. (2018) Naturally occurring variations in the Nod-independent model legume *Aeschynomene evenia* and relatives: a resource for nodulation genetics. BMC Plant Biology 18, 54.

Chen Y, Chen L, Lun ATL, Baldoni PL, Smyth GK (2024) edgeR 4.0: powerful differential analysis of sequencing data with expanded functionality and improved support for small counts and larger datasets. bioRxiv. 10.1101/2024.01.21.576131

Choi IS, Jansen R, Ruhlman T (2019) Lost and found: return of the inverted repeat in the legume clade defined by its absence. Genome Biology and Evolution 11, 1321–1333.

Danecek P, Bonfield JK, Liddle J, Marshall J, Ohan V, Pollard MO, Whitwham A, Keane T, McCarthy SA, Davies RM (2021) Twelve years of SAMtools and BCFtools. GigaScience. 10.1093/gigascience/giab008

De Carvalho-Niebel F, Fournier J, Becker A, Marín Arancibia M (2024) Cellular insights into legume root infection by rhizobia. Current Opinion in Plant Biology 81, 102597.

Djébali N, Jauneau A, Carine AT, Chardon F, Jaulneau V, Mathé C, Bottin A, Cazaux M, Pilet-Nayel ML, Baranger A et al. (2009) Partial resistance of *Medicago truncatula* to *Aphanomyces euteiches* is associated with protection of the root stele and is controlled by a major QTL rich in proteasome-related genes. Molecular Plant-Microbe Interactions 22, 1043–1055.

Djébali N, Mhadhbi H, Lafitte C, Dumas B, Esquerré-Tugayé MT, Aouani ME, Jacquet C (2011) Hydrogen peroxide scavenging mechanisms are components of *Medicago truncatula* partial resistance to *Aphanomyces euteiches*. European Journal of Plant Pathology 131, 559–571.

Dobin A, Davis CA, Schlesinger F, Drenkow J, Zaleski C, Jha S, Batut P, Chaisson M, Gingeras TR (2013) STAR: ultrafast universal RNA-seq aligner. Bioinformatics 29, 15–21.

Dubois M, Van den Broeck L, Inzé D (2018) The pivotal role of ethylene in plant growth. Trends in Plant Science 23, 311–323.

Emms DM, Kelly S (2019) OrthoFinder: phylogenetic orthology inference for comparative genomics. Genome Biology 20, 238.

Ewels PA, Peltzer A, Fillinger S, Patel H, Alneberg J, Wilm A, Garcia MU, Di Tommaso P, Nahnsen S (2020) The nf-core framework for community-curated bioinformatics pipelines. Nature Biotechnology 38, 276–278.

Feng Y, Wu P, Liu C, Peng L, Wang T, Wang C, Tan Q, Li B, Ou Y, Zhu H et al. (2021) Suppression of LjBAK1-mediated immunity by SymRK promotes rhizobial infection in *Lotus japonicus*. Molecular Plant 14, 1935–1950.

Gaulin E, Jacquet C, Bottin A, Dumas B (2007) Root rot disease of legumes caused by *Aphanomyces euteiches*. Molecular Plant Pathology 8, 539–548.

Gibelin-Viala C, Amblard E, Puech-Pages V, Bonhomme M, Garcia M, Bascaules-Bedin A, Fliegmann J, Wen J, Mysore KS, Le Signor C et al. (2019) The *Medicago truncatula* LysM receptor-like kinase LYK9 plays a dual role in immunity and arbuscular mycorrhizal symbiosis. New Phytologist 223, 1516–1522.

Gully D, Czernic P, Cruveiller S, Mahé F, Longin C, Vallenet D, François P, Nidelet S, Rialle S, Giraud E et al. (2018) Transcriptome profiles of Nod factor-independent symbiosis in the tropical legume *Aeschynomene evenia*. Scientific Reports 8, 10934.

Gu Z, Eils R, Schlesner M (2016) Complex heatmaps reveal patterns and correlations in multidimensional genomic data. Bioinformatics 32, 2847–2849.

Hailemariam S, Liao CJ, Mengiste T (2024) Receptor-like cytoplasmic kinases: orchestrating plant cellular communication. Trends in Plant Science 29, 432–445.

Herridge DF, Peoples MB, Boddey RM (2008) Global inputs of biological nitrogen fixation in agricultural systems. Plant and Soil 311, 1–18.

Horta-Araújo N, Landry D, Quilbé J, Pervent M, Nouwen N, Klopp C, Cullimore J, Gully D, Vicedo C, Gasciolli V et al. (2025) The receptor-like cytoplasmic kinase AeRLCK2 mediates Nod-independent rhizobial symbiosis in *Aeschynomene* legumes. The Plant Cell 37, 3456–3472.

Horta Araújo N, Nouwen N, Arrighi JF (2024) Nodulating another way: what can we learn from lateral root base nodulation in legumes? Journal of Experimental Botany 75, 3214–3219.

Johansson A, Sarrette B, Boscari A, Prudent M, Gruber V, Brouquisse R, Jacquet C, Gough C, Pauly N (2025) The role of reactive oxygen, nitrogen, and sulfur species in the integration of (a)biotic stress signals in legumes. Journal of Experimental Botany 76, 3774–3792.

Jones P, Binns D, Chang HY, Fraser M, Li W, McAnulla C, McWilliam H, Maslen J, Mitchell A, Nuka G, Pesseat S, Quinn AF, Sangrador-Vegas A, Scheremetjew M, Yong SY, Lopez R, Hunter S (2014) InterProScan 5: genome-scale protein function classification. Bioinformatics 30, 1236–1240.

Kälin C, Piombo E, Bourras S, Brantestam AK, Dubey M, Elfstrand M, Karlsson M (2024) Transcriptomic analysis identifies candidate genes for Aphanomyces root rot disease resistance in pea. BMC Plant Biology 24, article 000.

Kebede E (2021) Contribution, utilization, and improvement of legumes-driven biological nitrogen fixation in agricultural systems. Frontiers in Sustainable Food Systems 5, 767998.

Kiirika LM, Bergmann HF, Schikowsky C, Wimmer D, Korte J, Schmitz U, Niehaus K, Colditz F (2012) Silencing of the Rac1 GTPase MtROP9 in *Medicago truncatula* stimulates early mycorrhizal and oomycete root colonisations but negatively affects rhizobial infection. Plant Physiology 159, 501–516.

Kopylova E, Noé L, Touzet H (2012) SortMeRNA: fast and accurate filtering of ribosomal RNAs in metatranscriptomic data. Bioinformatics 28, 3211–3217.

Kovaka S, Zimin AV, Pertea GM, Razaghi R, Salzberg SL, Pertea M (2019) Transcriptome assembly from long-read RNA-seq alignments with StringTie2. Genome Biology 20, 278.

Krueger F, James F, Ewels P, Afyounian E, Schuster-Boeckler B (2021) TrimGalore v0.6.7. Available at: 10.5281/zenodo.5127899.

Kumar NTS, Caudillo-Ruiz KB, Chatterton S, Banniza S (2021) Characterization of *Aphanomyces euteiches* pathotypes infecting peas in Western Canada. Plant Disease 105, 10.1094.

Larsen LJ, Schlatter DC, Nimpoeno J, Hines-Snider C, Samac DA (2023) Rhizosphere and root community analysis of oomycetes associated with poor alfalfa (*Medicago sativa*) seedling establishment. Phytobiomes Journal 7, 526–537.

Lavaud C, Lesné A, Leprévost T, Pilet-Nayel ML (2024) Fine mapping of Ae-Ps4.5, a major locus for resistance to pathotype III of *Aphanomyces euteiches* in pea. Theoretical and Applied Genetics 137, 1–14.

Lavaud C, Lesné A, Piriou C, Roy GL, Boutet G, Moussart A, Poncet C, Delourme R, Baranger A, Pilet-Nayel ML (2015) Validation of QTL for resistance to *Aphanomyces euteiches* in different pea genetic backgrounds using near-isogenic lines. Theoretical and Applied Genetics 128, 2273–2288.

Li B, Dewey CN (2011) RSEM: accurate transcript quantification from RNA-Seq data with or without a reference genome. BMC Bioinformatics 12, 323.

Love MI, Huber W, Anders S (2014) Moderated estimation of fold change and dispersion for RNA-seq data with DESeq2. Genome Biology 15, 550.

Love MI, Soneson C, Hickey PF, Johnson LK, Pierce NT, Shepherd L, Morgan M, Patro R (2020) Tximeta: reference sequence checksums for provenance identification in RNA-seq. PLoS Computational Biology 16, e1007664.

Miyakawa T, Hatano KI, Miyauchi Y, Suwa YI, Sawano Y, Tanokura M (2014) A secreted protein with plant-specific cysteine-rich motif functions as a mannose-binding lectin exhibiting antifungal activity. Plant Physiology 166, 766–778.

Mou B, Zhao G, Wang J, Wang S, He F, Ning Y, Li D, Zheng X, Cui F, Xue F, et al. (2024) The OsCPK17–OsPUB12–OsRLCK176 module regulates immune homeostasis in rice. The Plant Cell 36, 987–1006.

Moussart A, Even MN, Tivoli B (2008) Reaction of genotypes from several grain and forage legumes to infection with a French pea isolate of *Aphanomyces euteiches*. European Journal of Plant Pathology 122, 321–333.

Okonechnikov K, Conesa A, García-Alcalde F (2016) Qualimap 2: advanced multi-sample quality control for high-throughput sequencing data. Bioinformatics 32, 292–294.

Patro R, Duggal G, Love MI, Irizarry RA, Kingsford C (2017) Salmon provides fast and bias-aware quantification of transcript expression. Nature Methods 14, 417–419.

Pertea G, Pertea M (2020) GFF utilities: GffRead and GffCompare. F1000Research 9, 304. doi:10.12688/f1000research.23297.2.

Pilet-Nayel ML, Muehlbauer FJ, McGee RJ, Kraft JM, Baranger A, Coyne CJ (2002) Quantitative trait loci for partial resistance to *Aphanomyces* root rot in pea. Theoretical and Applied Genetics 106, 28–39.

Quilbé J, Lamy L, Brottier L, Leleux P, Fardoux J, Rivallan R, Benichou T, Guyonnet R, Becana M, Villar I, et al. (2021) Genetics of nodulation in *Aeschynomene evenia* uncovers mechanisms of the rhizobium–legume symbiosis. Nature Communications 12, 1–14.

Quilbé J, Nouwen N, Pervent M, Guyonnet R, Cullimore J, Gressent F, Horta-Araújo N, Gully D, Klopp C, Giraud E, et al. (2022) A mutant-based analysis of the establishment of Nod-independent symbiosis in *Aeschynomene evenia*. Plant Physiology 190, 1400–1417.

Quinlan AR, Hall IM (2010) BEDTools: a flexible suite for comparing genomic features. Bioinformatics 26, 841–842.

Rey T, Jacquet C (2018) Symbiosis genes for immunity and vice versa. Current Opinion in Plant Biology 44, 64–71.

Rey T, Nars A, Bonhomme M, Bottin A, Huguet S, Balzergue S, Jardinaud MF, Bono JJ, Cullimore J, Dumas B, et al. (2013) NFP, a LysM protein controlling Nod factor perception, also intervenes in *Medicago truncatula* resistance to pathogens. New Phytologist 198, 875–886.

Roux F, Voisin D, Badet T, Balagué C, Barlet X, Huard-Chauveau C, Roby D, Raffaele S (2014) Resistance to phytopathogens e tutti quanti: placing plant quantitative disease resistance on the map. Molecular Plant Pathology 15, 427–436.

Sarrette B, Luu TB, Johansson A, Fliegmann J, Pouzet C, Pichereaux C, Remblière C, Sauviac L, Carles N, Amblard E, et al. (2025) *Medicago truncatula* SOBIR1 controls pathogen immunity and specificity in the Rhizobium–legume symbiosis. *Plant*, Cell & Environment 48, 32–50.

Savary S, Willocquet L, Pethybridge SJ, Esker P, McRoberts N, Nelson A (2019) The global burden of pathogens and pests on major food crops. Nature Ecology & Evolution 3, 430–439.

Sayols S, Scherzinger D, Klein H (2016) dupRadar: a Bioconductor package for assessing PCR artifacts in RNA-seq data. BMC Bioinformatics 17, 428.

Schneider CA, Rasband WS, Eliceiri KW (2012) NIH Image to ImageJ: 25 years of image analysis. Nature Methods 9, 671–675.

Sun L, Zhang J (2020) Regulatory role of receptor-like cytoplasmic kinases in early immune signalling in plants. FEMS Microbiology Reviews 44, 845–856.

Tang D, Chen M, Huang X, Zhang G, Zeng L, Zhang G, Wu S, Wang Y (2023) SRplot: a free online platform for data visualization and graphing. PLOS ONE 18, e0294236.

Tang D, Wang G, Zhou J-M (2017) Receptor kinases in plant–pathogen interactions: more than pattern recognition. The Plant Cell 29, 618–637.

Tommaso PD, Chatzou M, Floden EW, Barja PP, Palumbo E, Notredame C (2017) Nextflow enables reproducible computational workflows. Nature Biotechnology 35, 316–319.

Truong HN, Fournier C, Pateyron S, Paysant-Le Roux C, Gravot A, Clément G, Jeandroz S (2025) Pathogen-induced root glutamine concentration is a determinant of the *Medicago truncatula–Aphanomyces euteiches* interaction outcome. Planta 262, 8.

Tsyganova AV, Brewin NJ, Tsyganov VE (2021) Structure and development of the legume–rhizobial symbiotic interface in infection threads. Cells 10, 1085.

Van Rossum G, Drake FL (2009) The Python Language Reference Release 3.1.1. Python Software Foundation.

Wall L, Christiansen T, Orwant J (2000) Programming Perl. O’Reilly Media, Sebastopol, CA.

Wang L, Wang S, Li W (2012) RSeQC: quality control of RNA-seq experiments. Bioinformatics 28, 2184–2185.

Wang D, Jin R, Shi X, Guo H, Tan X, Zhao A, Lian X, Dai H, Li S, Xin K, Tian C, Yang J, Chen W, Macho AP, Wang E (2025) A kinase mediator of rhizobial symbiosis and immunity in *Medicago*. Nature 643, 768–775.

Waszczak C, Carmody M, Kangasjärvi J (2018) Reactive oxygen species in plant signalling. Annual Review of Plant Biology 69, 209–236.

Wickham H, François R, Henry L, Müller K, Vaughan D (2014) dplyr: a grammar of data manipulation. Available at: https://cran.r-project.org.

Wickham H (2016) ggplot2: Elegant Graphics for Data Analysis. Springer-Verlag New York.

Wickham H, Averick M, Bryan J, Chang W, D’ L, McGowan A, François R, Grolemund G, Hayes A, Henry L, et al. (2019) Welcome to the tidyverse. Journal of Open Source Software 4, 1686.

Wu T, Hu E, Xu S, Chen M, Guo P, Dai Z, Feng T, Zhou L, Tang W, Zhan L, et al. (2021) clusterProfiler 4.0: a universal enrichment tool for interpreting omics data. Innovation 2, 100141.

Yadav H, Dreher D, Athmer B, Porzel A, Gavrin A, Baldermann S, Tissier A, Hause B (2019) *Medicago* terpene synthase 10 is involved in defense against an oomycete root pathogen. Plant Physiology 180, 1598–1613.

Zhao Z, Liu H, Wang C, Xu JR (2014) Comparative analysis of fungal genomes reveals differences in plant cell wall–degrading capacity in fungi (Erratum). BMC Genomics 15, 6.

Zou S, Tang Y, Xu Y, Ji J, Lu Y, Wang H, Li Q, Tang D (2022) TuRLK1, a leucine-rich repeat RLK indispensable for stripe rust resistance of YrU1 and broad pathogen resistance. BMC Plant Biology 22, 1–13.

