## Supplemental Methods for "A novel pathosystem between *Aeschynomene evenia* and *Aphanomyces euteiches* reveals new immune components in quantitative legume root-rot resistance"

### 1. Cortex colonisation analysis

library(ggplot2)

library(data.table)

library(dplyr)

Pourcentage=as.data.frame(fread("Cortex_colonisation.csv",na.strings=c("",NA,"NULL"),dec="."))

Col_gg=c("#619CFF","#F8766D")

Pourcentage$Genotype <- factor(Pourcentage$Genotype, levels = c('M','G'))

#WT

ggplot(Pourcentage, aes(x = Condition, y = Percent_of_colonisation, fill = Condition, alpha = 0.5)) +

geom_boxplot(outlier.alpha = 0, alpha = 0.7, lwd = 0.7) +

geom_jitter(aes(color = Condition), size = 2, width = 0.2, alpha = 0.8) +

scale_fill_manual(values = Col_gg) +

scale_color_manual(values = Col_gg) +

stat_summary(fun = mean, geom = "point", shape = 23, size = 3, color = "black", fill = "black") +

facet_grid(~ Genotype, labeller = labeller(Genotype = as_labeller(c(

'M' = "n°76",

'G' = "n°79"

), default = label_value))) +

ylim(0, 5) +

ggtitle("") +

xlab("Condition") +

ylab("Percent of colonised cortex surface") +

scale_x_discrete(limits = c("ni", "RB84")) +

theme_classic(base_size = 14) +

theme(

strip.text = element_text(face = "bold", size = 12),

axis.text.x = element_text(angle = 0, vjust = 0.5),

strip.placement = "outside",

strip.background = element_blank(),

panel.spacing = unit(1, "lines")

)

\#### Stats

Non_Inoc<- Pourcentage[Pourcentage$Condition == "ni",]

Inoc <- Pourcentage[Pourcentage$Condition == "RB84",]

attach(Non_Inoc)

pairwise.wilcox.test(Percent_of_colonisation,Genotype, p.adjust.method="bonferroni")

attach(Inoc)

wilcox.test(Percent_of_colonisation[Genotype %in% c("M", "G")] ~ Genotype[Genotype %in% c("M", "G")])

\# Mutants

Pourcentage=as.data.frame(fread("Cortex_colonisation.csv",na.strings=c("",NA,"NULL"),dec="."))

Col_gg=c("#619CFF","#F8766D")

Pourcentage$Genotype <- factor(Pourcentage$Genotype, levels = c('M','X30','V21'))

ggplot(Pourcentage, aes(x = Condition, y = Percent_of_colonisation, fill = Condition, alpha = 0.5)) +

geom_boxplot(outlier.alpha = 0, alpha = 0.7, lwd = 0.7) +

geom_jitter(aes(color = Condition), size = 2, width = 0.2, alpha = 0.8) +

scale_fill_manual(values = Col_gg) +

scale_color_manual(values = Col_gg) +

stat_summary(fun = mean, geom = "point", shape = 23, size = 3, color = "black", fill = "black") +

facet_grid(~ Genotype, labeller = labeller(Genotype = as_labeller(c(

'M' = "n°76",

'X30' = "rlck2-1" ,

'V21' = "rlck2-2"

), default = label_value))) +

ylim(0, 5) +

ggtitle("") +

xlab("Condition") +

ylab("Percent of colonised cortex surface") +

scale_x_discrete(limits = c("ni", "RB84")) +

theme_classic(base_size = 14) +

theme(

strip.text = element_text(face = "bold", size = 12),

axis.text.x = element_text(angle = 0, vjust = 0.5),

strip.placement = "outside",

strip.background = element_blank(),

panel.spacing = unit(1, "lines")

)

\#### Stats

Non_Inoc<- Pourcentage[Pourcentage$Condition == "ni",]

Inoc <- Pourcentage[Pourcentage$Condition == "RB84",]

attach(Non_Inoc)

pairwise.wilcox.test(Percent_of_colonisation,Genotype, p.adjust.method="bonferroni")

attach(Inoc)

wilcox.test(Percent_of_colonisation[Genotype %in% c("M", "X30")] ~ Genotype[Genotype %in% c("M", "X30")])

wilcox.test(Percent_of_colonisation[Genotype %in% c("M", "V21")] ~ Genotype[Genotype %in% c("M", "V21")])

### 2. Phenotyping data, statistics and plots

library(ggplot2)

library(data.table)

library(dplyr)

############### WT ####################

Pourcentage=as.data.frame(fread("WT_ABCD.csv",na.strings=c("",NA,"NULL"),dec="."))

Inoc <- Pourcentage[Pourcentage$Condition == "RB84"| Pourcentage$Condition == "All2",]

Col=c("#FF3000","#75b8d1","#d18975", "#B276B2", "#60BD68" )

Col_gg=c("#00BA38","#619CFF","#F8766D")

\### Just M (n°76)

Pourcentage_M <- Pourcentage[Pourcentage$Genotype %in% c("M"), ]

Inoc_M <- Pourcentage_M[Pourcentage_M$Condition == "RB84"| Pourcentage_M$Condition == "All2",]

#Fresh Weight comparison

attach(Pourcentage_M)

pairwise.wilcox.test(Poids,Condition,p.adjust.method="bonferroni")

#Browning comparison

attach(Inoc_M)

pairwise.wilcox.test(Pourcentage_3dpi,Condition,p.adjust.method="bonferroni")

pairwise.wilcox.test(Pourcentage_5dpi,Condition,p.adjust.method="bonferroni")

pairwise.wilcox.test(Pourcentage_7dpi,Condition,p.adjust.method="bonferroni")

pairwise.wilcox.test(Pourcentage_10dpi,Condition,p.adjust.method="bonferroni")

pairwise.wilcox.test(Pourcentage_12dpi,Condition,p.adjust.method="bonferroni")

#Weight plot

ggplot(Pourcentage_M, aes(x = Condition, y = Poids, fill=Condition, alpha=0.5)) +

geom_boxplot(outlier.alpha = 0, alpha = 0.7, lwd = 0.7) +

geom_jitter(aes(color = Condition, shape = Rep), size = 2, width = 0.2, alpha = 0.8) +

scale_fill_manual(values = Col_gg) +

scale_color_manual(values = Col_gg) +

facet_grid(~ Genotype, labeller = labeller(Genotype = as_labeller(c(

'M' = "M" ), default = label_parsed))) +

stat_summary(fun = mean, geom = "point", shape = 23, size = 3, color = "black", fill = "black") +

ylim(0.02, 0.15) +

ggtitle("Fresh weight 21 dpi") +

xlab("Condition") +

ylab("Weight (g)") +

scale_x_discrete(limits = c("ni","All2", "RB84")) +

theme_classic(base_size = 14) +

theme(strip.text = element_text(face = "bold", size = 12))

#Kinetics of browning

Inoc_loop=Pourcentage_M %>% select(Condition, Pourcentage_3dpi, Pourcentage_5dpi, Pourcentage_7dpi, Pourcentage_10dpi, Pourcentage_12dpi)

day=3

mean_per_cond=Pourcentage_M %>% group_by(Condition) %>% summarise(moy_perc=mean(!!as.symbol(paste0("Pourcentage_",day,"dpi")), na.rm=T))

mean_per_cond$day=rep(day, nrow(mean_per_cond))

for (day in c(3,5,7,10,12)) {

#day=3

mean_per_cond_interm=Pourcentage_M %>% group_by(Condition) %>% summarise(moy_perc=mean(!!as.symbol(paste0("Pourcentage_",day,"dpi")), na.rm=T))

mean_per_cond_interm$day=rep(day, nrow(mean_per_cond_interm))

mean_per_cond=bind_rows(mean_per_cond, mean_per_cond_interm)

}

mean_per_cond$Condition <- recode(mean_per_cond$Condition, "All2" = "ATCC201834")

mean_per_cond <- data.frame()

for (day in c(3, 5, 7, 10, 12)) {

col_name <- paste0("Pourcentage_", day, "dpi")

mean_per_cond_interm <- Pourcentage_M %>%

group_by(Condition) %>%

summarise(

moy_perc = mean(!!as.symbol(col_name), na.rm = TRUE),

se_perc = sd(!!as.symbol(col_name), na.rm = TRUE) / sqrt(n())

)

mean_per_cond_interm$day <- day

mean_per_cond <- bind_rows(mean_per_cond, mean_per_cond_interm)

}

mean_per_cond$day <- as.numeric(mean_per_cond$day)

#Kinetics browning plot

ggplot(mean_per_cond, aes(x = day, y = moy_perc, color = Condition)) +

geom_point(size = 2) +

geom_line(size = 1) +

geom_errorbar(aes(ymin = moy_perc - se_perc, ymax = moy_perc + se_perc), width = 0.2) +

scale_color_manual(values = Col_gg) +

theme_light() +

scale_x_continuous(breaks = c(3, 5, 7, 10, 12)) +

ggtitle("Kinetics of Average Brown Root Surface") +

xlab("dpi") +

ylab("Percentage of Brown Surface Area") +

ylim(0, 80)+

theme_classic(base_size = 14) +

theme(strip.text = element_text(face = "bold", size = 12))

################ All natural accessions only RB84 (Rep A with Strain ATCC201834) ####################

Pourcentage_no_A <- Pourcentage %>% filter(!Rep %in% c("a"))

Inoc_no_A <- Pourcentage_no_A[Pourcentage_no_A$Condition == "RB84",]

#Weight comparison

attach(Pourcentage_no_A)

pairwise.wilcox.test(Poids,Geno_Cond,p.adjust.method="bonferroni")

#Browning comparison

attach(Pourcentage_no_A)

pairwise.wilcox.test(AUC_Brun, Genotype, p.adjust.method="bonferroni")

Pourcentage_no_A$Genotype <- factor(Pourcentage_no_A$Genotype, levels = c('M', '21', 'W', 'H', 'G'))

Inoc_no_A$Genotype <- factor(Inoc_no_A$Genotype, levels = c('M', '21', 'W', 'H', 'G'))

#Weight plot

ggplot(Pourcentage_no_A, aes(x = Condition, y = Poids, fill = Condition), alpha=0.5) +

geom_boxplot(outlier.alpha = 0, alpha = 0.7, lwd = 0.7) +

geom_jitter(aes(color = Condition, shape = Rep), size = 2, width = 0.2, alpha = 0.8) +

scale_fill_manual(values = Col_gg) +

scale_color_manual(values = Col_gg) +

facet_grid(~ Genotype, labeller = labeller(Genotype = as_labeller(c(

'M' = "M",

'21' = "21",

'W' = "W",

'H' = "H",

'G' = "G"

), default = label_parsed))) +

stat_summary(fun = mean, geom = "point", shape = 23, size = 3, color = "black", fill = "black") +

ylim(0.02, 0.15) +

ggtitle("") +

xlab("Condition") +

ylab("Fresh weight (g)") +

scale_x_discrete(limits = c("ni", "RB84")) +

theme_classic(base_size = 14) +

theme(strip.text = element_text(face = "bold", size = 12))

#Browning plot

ggplot(Inoc_no_A, aes(x = Genotype, y = AUC_Brun, fill = Genotype, alpha=0.5)) +

geom_boxplot(outlier.alpha = 0, alpha = 0.7, lwd = 0.7) +

geom_jitter(aes(color = Genotype, shape = Rep), size = 2, width = 0.2, alpha = 0.8) +

scale_fill_manual(values = Col) +

scale_color_manual(values = Col) +

stat_summary(fun = mean, geom = "point", shape = 23, size = 3, color = "black", fill = "black") +

ylim(0, 750) +

ggtitle("AUC of percentage of brown surface area") +

xlab("Genotype") +

ylab("AUC") +

scale_x_discrete(limits = c('M', '21','W','H','G'))+

theme_classic(base_size = 14) +

theme(

axis.text.x = element_text(angle = 0, vjust = 0.5),

strip.text = element_text(face = "bold", size = 12)

)

############### Mutants ###########################

Pourcentage=as.data.frame(fread("Mutants_Paper.csv",na.strings=c("",NA,"NULL"),dec="."))

Pourcentage_ <- Pourcentage[Pourcentage$Genotype %in% c("M", "V21", "X30", "E26", 'J42'), ]

Inoc_ <- Pourcentage_[Pourcentage_$Condition == "RB84",]

Pourcentage_$Poids <- as.numeric(Pourcentage_$Poids)

#Weight comparison

attach(Pourcentage_)

pairwise.wilcox.test(Poids,Geno_Cond,p.adjust.method="bonferroni")

#Check Replicate+Genotype interaction

fit_AUC = lm(AUC_Brun ~ Genotype*Rep, data = Inoc_, contrasts = list(Genotype = contr.treatment(5, 3))) # WT as reference

summary(fit_AUC)

#No interaction so linear model with replicate effect

fit_AUC = lm(AUC_Brun ~ Genotype + Rep, data = Inoc_, contrasts = list(Genotype = contr.treatment(5, 3))) # WT as reference

summary(fit_AUC)

Col=c("#FF3000","#0072B2","#56B4E9","#E69F00", "#FFCC00")

Col_gg=c("#619CFF","#F8766D")

Pourcentage_$Genotype <- factor(Pourcentage_$Genotype, levels = c('M','V21', 'X30', "E26", 'J42'))

Inoc_$Genotype <- factor(Inoc_$Genotype, levels = c('M','V21', 'X30', 'E26', 'J42'))

#Weight plot

ggplot(Pourcentage_, aes(x = Condition, y = Poids, fill = Condition, alpha = 0.5)) +

geom_boxplot(outlier.alpha = 0, alpha = 0.7, lwd = 0.7) +

geom_jitter(aes(color = Condition, shape = Rep), size = 2, width = 0.2, alpha = 0.8) +

scale_fill_manual(values = Col_gg) +

scale_color_manual(values = Col_gg) +

stat_summary(fun = mean, geom = "point", shape = 23, size = 3, color = "black", fill = "black") +

facet_grid(~ Genotype, labeller = labeller(Genotype = as_labeller(c(

'M' = "WT",

'V21' = "italic('rlck2-1')",

'X30' = "italic('rlck2-2')",

'E26' = "italic('crk-1')",

'J42' = "italic('crk-2')"), default = label_parsed))) +

ylim(0.03, 0.2) +

ggtitle("") +

xlab("Condition") +

ylab("Fresh weight (g)") +

scale_x_discrete(limits = c("ni", "RB84")) +

theme_classic(base_size = 14) +

theme(

strip.text = element_text(face = "bold", size = 12),

axis.text.x = element_text(angle = 0, vjust = 0.5),

strip.placement = "outside",

strip.background = element_blank(),

panel.spacing = unit(1, "lines")

)

#Browning plot

ggplot(Inoc_, aes(x = Genotype, y = AUC_Brun, fill = Genotype, alpha=0.5)) +

geom_boxplot(outlier.alpha = 0, alpha = 0.7, lwd = 0.7) +

geom_jitter(aes(color = Genotype, shape = Rep), size = 2, width = 0.2, alpha = 0.8) +

scale_fill_manual(values = Col) +

scale_color_manual(values = Col) +

stat_summary(fun = mean, geom = "point", shape = 23, size = 2, color = "black", fill = "black") +

ylim(0, 650) +

ggtitle("") +

xlab("Genotype") +

ylab("AUC of brown surface area") +

scale_x_discrete(labels = c(

'V21' = expression(italic("rlck2-1")),

'X30' = expression(italic("rlck2-2")),

'E26' = expression(italic("crk-1")),

'J42' = expression(italic("crk-2"))

)) + theme_classic(base_size = 14) +

theme(

axis.text.x = element_text(angle = 0, vjust = 0.5),

strip.text = element_text(face = "bold", size = 12)

)

### 3. DEG enrichment analysis template

\# Perform functional enrichment analysis

\# Pipeline: clusterprofiler, GO analysis for non-model organisms

\# Modified from: https://yulab-smu.top/biomedical-knowledge-mining-book/index.html

\# if (!require("BiocManager", quietly = TRUE))

\#   install.packages("BiocManager")

\# BiocManager::install("clusterProfiler")

library(clusterProfiler)

library(ggplot2)

library(dplyr)

library(tidyr)

library(edgeR)

library(ape)

library(stringr)

library(gsubfn)

rm(list=ls())

setwd("\\\\BIO66\\evo\\commun\\projects\\patho_evo\\aphano\\pea_vs_aphano\\degs\\pi_vs_strain16\\48h")

#########################################

\## Part 1: GO terms

#########################################

\# -----------------------

\## 1. Build a database of with user-defined gene and term annotations

\# -----------------------

\# Motivation: clustalprofiler and other enrichment analysis provide easy resources only for model organisms (and sometimes an outdated genome version, or with a different gene ids) -> solution: work with user-defined gene and term annotations

\## Prepare data frame with all gene ids, go ids and go terms

\# Import interproscan annotations

\# NOTE: This is the interproscan output fixed to have the same number of columns in all rows, for that:

\# awk -F "\t" 'BEGIN {OFS="\t"} {if (NF == 13) $14 = "-"; print}' Marpal.tsv > Marpal_equal_columns.tsv

ips <- read.csv("Interpro_equal_columns.tsv", h = F, sep = "\t", na.strings = c("", "-", "NA"))

colnames(ips) <- c("Protein_accession","Sequence_MD5","Sequence_length",

                   "Analysis","Signature_accession","Signature_description",

                   "Start_location","Stop_location","Score","Status","Date",

                   "InterPro_accession","InterPro_description",

                   "GO_annotations")

head(ips)

dim(ips)

length(unique(ips$Protein_accession))

ips <- ips %>% mutate(Protein_accession = gsub("\\.\\d+\\b", "", ips$Protein_accession))

\# Keep only gene id and go id

ips.go <- ips[,c("Protein_accession", "GO_annotations")]

head(ips.go)

\# Remove NAs

dim(ips.go)

if(any(is.na(ips.go$GO_annotations))){

  ips.go <- ips.go[is.na(ips.go$GO_annotations) == F,]

}

dim(ips.go)

head(ips.go)

\# Split entries with multiple GO separated by "|", and remove duplicates

ips.go <- ips.go %>%

  separate_rows(GO_annotations, sep = "\\|") %>%

  distinct()

head(ips.go)

\# Extract GO terms

goterms <- go2term(ips.go$GO_annotations)

colnames(goterms)[1] <- colnames(ips.go)[2] # Match colnames so it can be more easily joined

head(goterms)

\# Add GO terms to original table

ips.go.complete <- ips.go %>%

  left_join(goterms, by = "GO_annotations")

head(ips.go)

head(ips.go.complete)

\# -----------------------

\## 2. Define lists of genes (background + up/down DEs)

\# -----------------------

\## Define set of background genes

analyzed_genes <- read.csv('degs_results', sep = '\t', header = T)

gff <- read.gff('Species_annotation_file.gff', GFF3 = T)

genes <- gff %>% filter(type == 'gene')

\# Change according species (Pissat = Pisum sativum, Aeseve for Aeschynomene evenia)

genes <- genes %>% mutate(gene_code = gsub("ID=|;", "", str_extract(attributes, "ID=Pissat_.{1,}?;")))

background_genes <- genes$gene_code

length(background_genes)

\# Downregulated

down_degs <- analyzed_genes %>% filter((status == 'down') & (logFC <= -1))

down_degs <- down_degs$genes

length(down_degs)

\# Upregulated

up_degs <- analyzed_genes %>% filter((status == 'up') & (logFC >= 1))

up_degs <- up_degs$genes

length(up_degs)

\# -----------------------

\## 3. Perform enrichment analysis (GO terms)

\# -----------------------

\## 3.1. Downregulated genes

\# name the condition for output file

down_name = ''

head(ips.go.complete)

enrich_down <- enricher(down_degs,

                        TERM2GENE = ips.go.complete[,c(3,1)],

                        TERM2NAME = ips.go.complete[,c(3,2)],

                        pvalueCutoff = 0.05,

                        universe = background_genes,

                        qvalueCutoff = 0.05)

enrich_down

write.table(enrich_down@result, paste("table_enrichment_GO_results", down_name, "down.tsv", sep = "_"),

            col.names = T, row.names = F, quote = F, sep = "\t")

\# Generate figures

if(any(enrich_down@result$p.adjust <= 0.05)){

  enrich_down@result$Description <- enrich_down@result$ID

  p <- dotplot(enrich_down,

               x= "geneRatio", # Options: GeneRatio, BgRatio, pvalue, p.adjust, qvalue

               color="p.adjust",

               orderBy = "x", # Options: GeneRatio, BgRatio, pvalue, p.adjust, qvalue

               showCategory=100,

               font.size=8) +

    ggtitle("dotplot for GO ORA")

  ggsave(filename = paste("figure_enrichment_GO_dotplot", down_name, "down.pdf", sep = "_"),

         plot =  p,  dpi = 300, width = 18, height = 18, units = "cm")

}

\## 3.2. Upregulated genes

up_name = ""

head(ips.go.complete)

enrich_up <- enricher(up_degs,

                        TERM2GENE = ips.go.complete[,c(3,1)],

                        TERM2NAME = ips.go.complete[,c(3,2)],

                        pvalueCutoff = 0.05,

                        universe = background_genes,

                        qvalueCutoff = 0.05)

enrich_up

write.table(enrich_up@result, paste("table_enrichment_GO_results", up_name, "up.tsv", sep="_"),

            col.names = T, row.names = F, quote = F, sep = "\t")

\# Generate figures

if(any(enrich_up@result$p.adjust <= 0.05)){

  enrich_up@result$Description <- enrich_up@result$ID

  p <- dotplot(enrich_up,

               x= "geneRatio", # Options: GeneRatio, BgRatio, pvalue, p.adjust, qvalue

               color="p.adjust",

               orderBy = "x", # Options: GeneRatio, BgRatio, pvalue, p.adjust, qvalue

               showCategory=100,

               font.size=8) +

    ggtitle("dotplot for GO ORA")

  ggsave(filename = paste("figure_enrichment_GO_dotplot", up_name, "up.pdf", sep = "_"),

         plot =  p,  dpi = 300, width = 18, height = 18, units = "cm")

}

#########################################

\## Part 2: Interpro annotations

#########################################

\# -----------------------

\## 1. Prepare data frame with all gene ids and IPR ids

\# -----------------------

\# Keep only HOG id, IPR accession and description

colnames(ips)

ips.interpro <- ips[,c(1,12)]

head(ips.interpro)

\# Remove NAs

dim(ips.interpro)

if(any(is.na(ips.interpro$InterPro_accession))){

  ips.interpro <- ips.interpro[is.na(ips.interpro$InterPro_accession) == F,]

}

dim(ips.interpro)

head(ips.interpro)

\# Split entries with multiple ipr separated by "|", and remove duplicates

ips.interpro <- ips.interpro %>%

  separate_rows(InterPro_accession, sep = "\\|") %>%

  distinct()

head(ips.interpro)

dim(ips.interpro)

\# Import IPR entry same version as InterProScan used

ips.entry <- read.csv("entry.list", h=T, sep="\t")

head(ips.entry)

ips.entry <- ips.entry[,c(1,3)]

colnames(ips.interpro)

colnames(ips.entry)

colnames(ips.entry) <- c(colnames(ips.interpro)[2], "InterPro_Description") # Match colnames to make it easier to join

head(ips.entry)

\# Add to final annotation table

ips.interpro <- ips.interpro %>%

  left_join(ips.entry, by = "InterPro_accession")

ips.interpro

\# -----------------------

\## 2. Perform enrichment analysis

\# -----------------------

\## 3.1. Downregulated genes

head(ips.interpro)

enrich_down <- enricher(down_degs,

                        TERM2GENE = ips.interpro[,c(3,1)],

                        TERM2NAME = ips.interpro[,c(3,2)],

                        pvalueCutoff = 0.05,

                        universe = background_genes,

                        qvalueCutoff = 0.05)

enrich_down

write.table(enrich_down@result, paste("table_enrichment_InterPro_results", down_name, "down.tsv", sep = "_"),

            col.names = T, row.names = F, quote = F, sep = "\t")

\# Generate figures

if(any(enrich_down@result$p.adjust <= 0.05)){

  enrich_down@result$Description <- enrich_down@result$ID

  p <- dotplot(enrich_down,

               x= "geneRatio", # Options: GeneRatio, BgRatio, pvalue, p.adjust, qvalue

               color="p.adjust",

               orderBy = "x", # Options: GeneRatio, BgRatio, pvalue, p.adjust, qvalue

               showCategory=100,

               font.size=8) +

    ggtitle("dotplot for InterPro ORA")

  ggsave(filename = paste("figure_enrichment_InterPro_dotplot",down_name,"down.pdf", sep = "_"),

         plot =  p,  dpi = 300, width = 21, height = 30, units = "cm")

}

\## 3.2. Upregulated genes

head(ips.interpro)

enrich_up <- enricher(up_degs,

                      TERM2GENE = ips.interpro[,c(3,1)],

                      TERM2NAME = ips.interpro[,c(3,2)],

                      pvalueCutoff = 0.05,

                      universe = background_genes,

                      qvalueCutoff = 0.05)

enrich_up

write.table(enrich_up@result, paste("table_enrichment_InterPro_results", up_name, "up.tsv", sep = "_"),

            col.names = T, row.names = F, quote = F, sep = "\t")

\# Generate figures

if(any(enrich_up@result$p.adjust <= 0.05)){

  enrich_up@result$Description <- enrich_up@result$ID

  p <- dotplot(enrich_up,

               x= "geneRatio", # Options: GeneRatio, BgRatio, pvalue, p.adjust, qvalue

               color="p.adjust",

               orderBy = "x", # Options: GeneRatio, BgRatio, pvalue, p.adjust, qvalue

               showCategory=100,

               font.size=8) +

    ggtitle("dotplot for InterPro ORA")

  ggsave(filename = paste("figure_enrichment_InterPro_dotplot", up_name, "up.pdf", sep = "_"),

         plot =  p,  dpi = 300, width = 21, height = 30, units = "cm")

}

**Script for filtering low- variance genes, generating of MDS plot, estimating DEGs and creating Volcano plots.**

library('edgeR')

library('ggplot2')

library('ggrepel')

library('dplyr')

library('tibble')

library('Glimma')

setwd("C:\\Users\\jean.keller\\Nextcloud\\pissat_x_aphano")

\# Loading data

metadata <- read.delim("pissat_x_aphano_metadata.tsv", sep = '\t', header = T, colClasses = "character")

all_counts <- read.delim("pissat_vs_aphano_allcounts.length_scaled.tsv", sep = '\t', header = T)

all_counts <- all_counts %>% select(-gene_name)

\# metadata$group <- paste(metadata$condition, metadata$time, sep = '.')

#################################################################################

##################### 1 hour

\# Filtering conditions

cond_name <- 'pi_st16_48h'

exp_metadata <- metadata %>% filter(((treatment == 'mock') | (treatment == 'high_virulence')) & (timepoint == "48h") & (genotype == "resistant"))

\# exp_metadata <- metadata

exp_counts <- all_counts %>%  select('gene_id', as.vector(exp_metadata$sample))

\# exp_counts <- all_counts

write.table(exp_counts, paste(cond_name, 'counts.tsv', sep = "_"), sep = '\t', quote = F, row.names = F)

\# Preparing matrix

exp_group <- relevel(as.factor(exp_metadata$treatment), ref = 'mock')

y <- DGEList(counts = exp_counts[,2:length(exp_counts)], genes = exp_counts[,1], group = exp_group)

levels(y$samples$group)

y$samples

design <- model.matrix(~exp_group)

\# Filtering low variable genes

keep <- filterByExpr(y)

table(keep)

y <- y[keep, , keep.lib.sizes = F]

y$samples

\# Normalizing data

y <- normLibSizes(y)

y$samples

y <- estimateDisp(y, design = design, robust = T)

y$common.dispersio

pdf(paste(cond_name, 'plotBCV.pdf', sep = "_"))

plotBCV(y)

dev.off()

p <- plotMDS(y)

d <- data.frame(p$x, p$y)

row.names(d) <- colnames(exp_counts[,-1])

d <- left_join(rownames_to_column(d), exp_metadata, by=c("rowname" = "sample"))

row.names(d) <- colnames(exp_counts[,-1])

pdf(paste(cond_name, 'plotMDS.pdf', sep = "_"))

ggplot(d, aes(p.x, p.y, label=rowname, color=treatment))+

  geom_point(size=2)+

  coord_cartesian(clip="off")+

  # scale_shape_manual(values=c(23,24, 25))+

  scale_color_manual(values=c("blue", "red"))+

  geom_text_repel(box.padding = 0.3, min.segment.length = 0, seed = 42, show.legend = F, max.overlaps = 20)+

  xlab(paste0("Leading logFC dim 1 (", round(p$var.explained[[1]]*100, 1), "%)"))+

  ylab(paste0("Leading logFC dim 2 (", round(p$var.explained[[2]]*100, 1), "%)"))

dev.off()

\# Estimating DEGs

fit <- glmQLFit(y, design)

qlf.treat.ctrl <- glmQLFTest(fit, coef = 2)

res.qlf <- topTags(qlf.treat.ctrl, n = Inf)

summary(decideTests(qlf.treat.ctrl))

qlf.degs <- decideTests(qlf.treat.ctrl)

y_ordered <- y$counts

rownames(y_ordered) <- y$genes$genes

y_ordered <- y_ordered[match(res.qlf$table$genes, rownames(y_ordered)),]

res.qlf$table$status <- ifelse(res.qlf$table$FDR <= 0.05 & res.qlf$table$logFC > 0, 1,

                               ifelse(res.qlf$table$FDR <= 0.05 & res.qlf$table$logFC < 0, -1, 0))

df.qlf <- data.frame(GeneID=res.qlf$table$genes, row.names=row.names(y_ordered))

glXYPlot(res.qlf$table$logFC, -log10(res.qlf$table$FDR), status=res.qlf$table$status, anno = df.qlf,

         side.main = 'GeneID', groups = exp_group, counts = y_ordered, xlab = 'logFC', ylab = '-log10(FDR)',

         cols = c("indianred2", "grey", "steelblue2"), html = paste(cond_name, 'interactive_volcano', sep = '_'))

res.qlf$table <- res.qlf$table %>% mutate(status = recode(as.character(status), '1' = 'up', '-1'='down', '0'='ns'))

write.table(res.qlf, paste(cond_name, 'qlf_res_all.degs.tsv', sep = '_'), sep = '\t', quote = F, row.names = F)

\# Volcano plots

pdf(paste(cond_name, 'qlf.volcano.pdf', sep = "_"))

plotMD(qlf.treat.ctrl)

abline(h=c(1, -1), col = "grey", lwd=3, lty=2)

dev.off()

save.image(paste0(cond_name, '.RData'))

### 4. Heatmap

library('dplyr')

library('reshape2')

library('ggplot2')

library('readr')

library('ComplexHeatmap')

library('circlize')

aeseve_symb_patho_logFC1 <- read_tsv("Aeseve_symb_patho_logFC1_for_heatmap.tsv", show_col_types = FALSE)

#Sum of LogFC

df <- aeseve_symb_patho_logFC1 %>% rowwise() %>% mutate(symb_sum = sum(c_across(starts_with('aeseve_symb')))) %>%

mutate(patho_sum = sum(c_across(starts_with('aeseve_x_aphano'))))

df <- df[, c(

"geneCode",

"aeseve_x_aphano_1dpi_qlf_res_all",

"aeseve_x_aphano_3dpi_qlf_res_all",

"aeseve_symb_144hpi_qlf_res_all",

"aeseve_symb_96hpi_qlf_res_all",

"aeseve_symb_48hpi_qlf_res_all",

"aeseve_symb_24hpi_qlf_res_all",

"aeseve_symb_6hpi_qlf_res_all",

"symb_sum",

"patho_sum")]

#Filter genes DEG in both symb & imm

df_coexp <- df %>% filter((symb_sum != 0) & (patho_sum != 0))

df_patho_symb <- df_coexp %>% select(-c(patho_sum, symb_sum))#remove sum colums

df_patho_symb <- as.data.frame(df_patho_symb)#check that dataframe

row.names(df_patho_symb) <- df_patho_symb[,1]#sets Gene code as row names

df_patho_symb <- df_patho_symb %>% select(-geneCode)#remove gene code column

df_patho_symb <- as.matrix(df_patho_symb)#makes matrix

Heatmap(df_patho_symb, clustering_method_rows='single', column_names_rot = 45, show_row_names = F, row_dend_width = unit(15, "mm"), cluster_columns = FALSE)

#Cluster cols TRUE if you want the conditions clustered False if you want order of table

#Heatmap with categories at end because need to id the mixed profile genes first

########### Filter for mixed profile genes ############

#make table with just symb data

symb_cols <- c("geneCode","aeseve_symb_144hpi_qlf_res_all",

"aeseve_symb_96hpi_qlf_res_all",

"aeseve_symb_48hpi_qlf_res_all",

"aeseve_symb_24hpi_qlf_res_all",

"aeseve_symb_6hpi_qlf_res_all")

symb_data <- aeseve_symb_patho_logFC1[, symb_cols, drop=FALSE]

#make table with just patho data

patho_cols <- c("geneCode",

"aeseve_x_aphano_1dpi_qlf_res_all",

"aeseve_x_aphano_3dpi_qlf_res_all")

patho_data <- aeseve_symb_patho_logFC1[, patho_cols, drop=FALSE]

\# dire dans quelles colonnes il faut chercher si il y a des > & < de 0 = gene with mixed expression prifle

symb_cols_LogFC<- c(

"aeseve_symb_144hpi_qlf_res_all",

"aeseve_symb_96hpi_qlf_res_all",

"aeseve_symb_48hpi_qlf_res_all",

"aeseve_symb_24hpi_qlf_res_all",

"aeseve_symb_6hpi_qlf_res_all")

patho_cols_LogFC<- c("aeseve_x_aphano_1dpi_qlf_res_all",

"aeseve_x_aphano_3dpi_qlf_res_all")

\# list line number with mixed profile

genes_mixed_symb <- row.names(symb_data)[

apply(symb_data[, symb_cols_LogFC], 1, function(x) any(x > 0) & any(x < 0))]

length(genes_mixed_symb)

genes_mixed_patho <- row.names(patho_data)[

apply(patho_data[, patho_cols_LogFC], 1, function(x) any(x > 0) & any(x < 0))]

length(genes_mixed_patho)

##Remove rows with mixed profile genes from the list of all DEGs

aeseve_symb_patho_logFC1_filtered <- aeseve_symb_patho_logFC1[

!row.names(aeseve_symb_patho_logFC1) %in% genes_mixed_symb, , drop = FALSE]

#Opposite profile genes

#Make a table with logFC sum but with only the genes without a mixed profile

sum_filtered <- aeseve_symb_patho_logFC1_filtered %>% rowwise() %>% mutate(symb_sum = sum(c_across(starts_with('aeseve_symb')))) %>%

mutate(patho_sum = sum(c_across(starts_with('aeseve_x_aphano'))))

#From the genes without a mixed profile makes new tables of opposite genes

up_symb_down_patho<- sum_filtered %>% filter((symb_sum>0) & (patho_sum<0))

down_symb_up_patho<- sum_filtered %>% filter((symb_sum<0) & (patho_sum>0))

down_symb_patho_commun<- sum_filtered %>% filter((symb_sum<0) & (patho_sum<0))

up_symb_patho_commun<- sum_filtered %>% filter((symb_sum>0) & (patho_sum>0))

####TABLES###

write.table(up_symb_down_patho,file = "DEGs_Up_symb_down_patho.tsv",sep = "\t", quote = F, row.names = F)

write.table(down_symb_up_patho,file = "DEGs_down_symb_up_patho.tsv",sep = "\t", quote = F, row.names = F)

write.table(up_symb_patho_commun,file = "DEGs_Up_symb_patho_commun.tsv",sep = "\t", quote = F, row.names = F)

write.table(down_symb_patho_commun,file = "DEGs_down_symb_patho_commun.tsv",sep = "\t", quote = F, row.names = F)

########## Heatmap with categories ##################

#Assign categories to all DEGs (df) and then filter to only keep DEGs that are coexpressed (df_patho_symb)

gene_categories <- df %>%

mutate(Category = case_when(

symb_sum > 0 & patho_sum > 0 ~ "Common Up",

symb_sum < 0 & patho_sum < 0 ~ "Common Down",

symb_sum > 0 & patho_sum < 0 ~ "Up symb / Down patho",

symb_sum < 0 & patho_sum > 0 ~ "Down symb / Up patho",

TRUE ~ "Other"

)) %>%

filter(geneCode %in% rownames(df_patho_symb)) %>%

select(geneCode, Category)

#List of genes codes of genes with mixed profiles (same as above but gives genes code instead for row number)

genes_mixed_symb_codes <- symb_data$geneCode[

apply(symb_data[, symb_cols_LogFC], 1, function(x) any(x > 0) & any(x < 0))]

length(genes_mixed_symb_codes)

#assign "other" category to genes with mixed profile

gene_categories <- gene_categories %>%

mutate(Category = ifelse(geneCode %in% genes_mixed_symb_codes, "Other", Category))

#check numer of common and opposit DEGs

gene_categories %>% count(Category)

\# Convert to named vector for annotation

gene_category_vector <- gene_categories$Category

names(gene_category_vector) <- gene_categories$geneCode

category_colors <- c(

"Common Up" = "#D55E00",

"Common Down" = "#619CFF",

"Up symb / Down patho" = "#CC79A7",

"Down symb / Up patho" = "#92C5DE",

"Other" = "#000000")

row_annot <- rowAnnotation(

Category = gene_category_vector,

col = list(Category = category_colors),

width = unit(5, "mm"))

#Heatmap with categories

Heatmap(

df_patho_symb,

clustering_method_rows = 'single',

column_names_rot = 45,

show_row_names = FALSE,

row_dend_width = unit(15, "mm"),

cluster_columns = TRUE,

right_annotation = row_annot

)
